# Machine-learning convergent melanocytic morphology despite noisy archival slides

**DOI:** 10.1101/2024.09.12.612732

**Authors:** M Tada, G Gaskins, S Ghandian, N Mew, MJ Keiser, ES Keiser

## Abstract

Melanocytic atypia, ranging from benign to malignant, often leads to diagnostic discordance, complicating its prediction by machine learning models. To overcome this, we paired H&E-stained histology images with contiguous or serial sections immunohistochemically (IHC) stained for melanocytic cells via antibodies for MelanA, MelPro, or SOX10. We developed a deep-learning pipeline to identify melanocytic atypia by digitizing a real-world archival dataset of 122 paired whole slide images from 61 confirmed melanoma in situ (MIS) cases at two institutions. Only 37.7% of the cases contained tissue pairs that matched well enough for deep learning. Nonetheless, the MelanA+MelPro models achieved an average area under the receiver-operating characteristic (AUROC) of 0.948 and an average area under the precision-recall curve (AUPRC) of 0.611, while the SOX10 models had an average of 0.867 AUROC and 0.433 AUPRC. Despite learning from biologically different IHC stains, the convolutional neural network (CNN) models independently exhibited an intuitive convergent rationale by explainable AI saliency calculations. Different antibodies, with nuclear versus cytoplasmic staining, provided complementary yet consistent information, which the CNNs integrated effectively. The resulting multi-antibody virtual stains identified morphologic cytologic and small-scale architectural features directly from H&E-stained histology images, which can assist pathologists in assessing cutaneous MIS.

## Introduction

Whole slide scanned formalin-fixed paraffin-embedded (FFPE) tissue image analysis was only accepted in a research context^1–8^ before the US Federal Drug Administration (FDA) approved the first whole slide imaging system for histopathology primary diagnosis in 2017^9^.

The first FDA-approved machine learning-enabled medical device was also in the field of anatomic pathology for automated assessment of cervical cytology slides in 1995^10^, but radiology and several other medical subspecialties quickly surpassed pathology in the number of FDA-approved machine learning-enabled medical devices^11^. Currently, fewer than 1% of the 882 FDA-approved devices are specific to FFPE tissue pathology, of which a single device is FDA-approved for automated histomorphologic organ-specific analysis^12^. Here, we develop an approach to making FFPE skin tissue histomorphologic machine-learning algorithms for researchers and practicing dermatopathologists.

Convolutional neural networks^13^ effectively identify and diagnose a range of pathologies^14^, including skin disease^15–17^, with diagnostic performance approaching that of a general practice pathologist in some cases^18,19^. Most of these achievements rely on a paradigm of supervised learning, in which networks learn from a corpus of well-labeled, human-curated images. These models can generate predicted virtual stains meant to approximately replicate particular immunohistochemical (IHC) stains directly from hematoxylin and eosin (H&E) images^20–22^. Although new techniques are emerging to relax the resolution of labeling required^15,23,24^, pathologist-labeled images remain the currently accepted computational “ground truth,” synonymous with the clinical “gold standard.” However, if there is uncertainty regarding histopathologic ground-truth labels, the classification task becomes muddled, and the model becomes less generalizable.

Melanocytic atypia is a particularly challenging field of pathology, with a spectrum of melanocytic atypia seen in benign (nevus), atypical (dysplastic nevus), and malignant (melanoma) lesions. Cytologically, melanocytes show a range of atypia, sometimes mimicking benign epidermal keratinocytes. Architecturally, atypical melanocytes display anywhere from benign-appearing to highly atypical growth patterns. Thus, pathologist interobserver agreement for melanocytic atypia in standard (H&E) histology images ranges between 33-68%, with atypical nevus versus malignant melanoma cases accounting for much of the diagnostic discordance^25,26^.

Various conventional classification methods for melanocytic atypia exist, with diagnostic top-line terminology ranging from eponymous labels to descriptive diagnoses with up to nine permutations of architectural and cytologic atypia. Currently, accepted diagnostic terminology includes “melanocytic atypia with mild/moderate/severe architectural disorder and mild/moderate/severe cytologic atypia,” which some diagnostic laboratories have shortened to “mild/moderate/severe dysplastic nevus,” or rarely just “dysplastic nevus,” “atypical nevus,” or “Clark’s nevus,” with a comment recommending surveillance or excision^27,28^. More recently, the World Health Organization (WHO) proposed a two-grade system for dysplastic nevi with size criteria and intermittently (“low-CSD”; cumulative sun damage) versus chronically (“high-CSD”) sun-exposed diagnostic pathways for melanoma classification^29^. At times, it is not possible to differentiate between a moderately dysplastic nevus versus one that requires excision due to incomplete biopsy or unknown therapeutic history. Such cases often receive the diagnosis “atypical melanocytic neoplasm” or “atypical melanocytic proliferation” to communicate the need for additional clinical correlation, including further sampling or excision.

Severely dysplastic nevi and melanoma in situ undergo similar surgical treatment, but the diagnoses impart crucial “atypical” versus “malignant/ cancer” implications to the patient. While researchers could collapse these two diagnoses into a single category for a supervised learning task, it would be clinically misleading. In supervised machine learning, the utility of a learned function is dependent upon the accuracy and reliability of the labels used to train it. To circumvent the issue of diagnostic discordance negatively contributing to our “ground-truth” labels, we apply a pathologist-agnostic method to identify melanocytic atypia in patient tissue sections.

Pathologists employ immunohistochemical stains to highlight specific cell types in diagnostically challenging cases. Correspondingly, our computational method uses the melanocytic immunohistochemical (IHC) stained sections from archival patient cases to auto-label (notate) approximate melanocyte location within corresponding H&E tissue sections of diagnosed cases of “melanoma in situ.” A convolutional neural network (CNN) ingests 64x64 micron tiles (high-power views) of image data from the H&E section for training, with the tile labels extracted from the paired but unseen IHC section as the source of the melanocyte-specific labeling. While IHC stains highlight specific cells, the field of view visible in a “tile” in these models is intentionally larger than the size of a single cell. Thus, a computational algorithm requires a binary “tile-level label” of “*increased* melanocytes present” vs. “*few to no* melanocytes present,” which is determined by a fixed stain-specific cutoff within each IHC-stain model. Just as pathologists do not study and diagnose melanoma by solely viewing a melanocytic immunohistochemical stain without an H&E paired section, we do not train our CNN on IHC-tissue samples directly. Instead, we assemble a custom dataset composed of adjacently sliced and paired tissue sections. Each set of paired tissue consists of at least one tissue level that is IHC-stained and a complementary level that is H&E-stained. We refer to both tissue profiles collectively as a “sample pair.”

The computational extraction of IHC-stain information for each sample pair allows the method to incorporate information regarding melanocyte location and morphology (Figure 1) while only training on H&E data with known pathologist-assigned diagnoses, e.g., melanoma in situ in this study. Ideally, the CNN learns features indicative of melanocytic atypia that are generalizable to new H&E images. While others have also adopted this paired-stain deep learning training strategy, their approach required laborious custom tissue restaining or creation of new “back to back” H&E and IHC paired slides instead of leveraging existing clinical archival slides of nonidentical tissue sections^22,30^ The method relies on a critical assumption: that sequential tissue sections from the same tissue block (some “contiguously” cut “back to back” by the same histotechnologist at the same point in time, and others cut “serially” or “discontinuously” at two different points in time with the tissue block re-faced for the IHC stain) are similar enough in morphology and location that IHC staining from one slice can serve as a proxy label for its adjacent (H&E) pair. To maximize this assumption and to ensure optimal morphological congruence among paired H&E and IHC images, we align sample pairs at their native 40x resolution. We then train a convolutional neural network (CNN), specifically DenseNet121,^31^ to identify melanocytic atypia in H&E images at the tile level. We refer to melanocytes within melanoma in situ as “atypical melanocytes” due to their association with malignant lesions.

**Figure 1.**
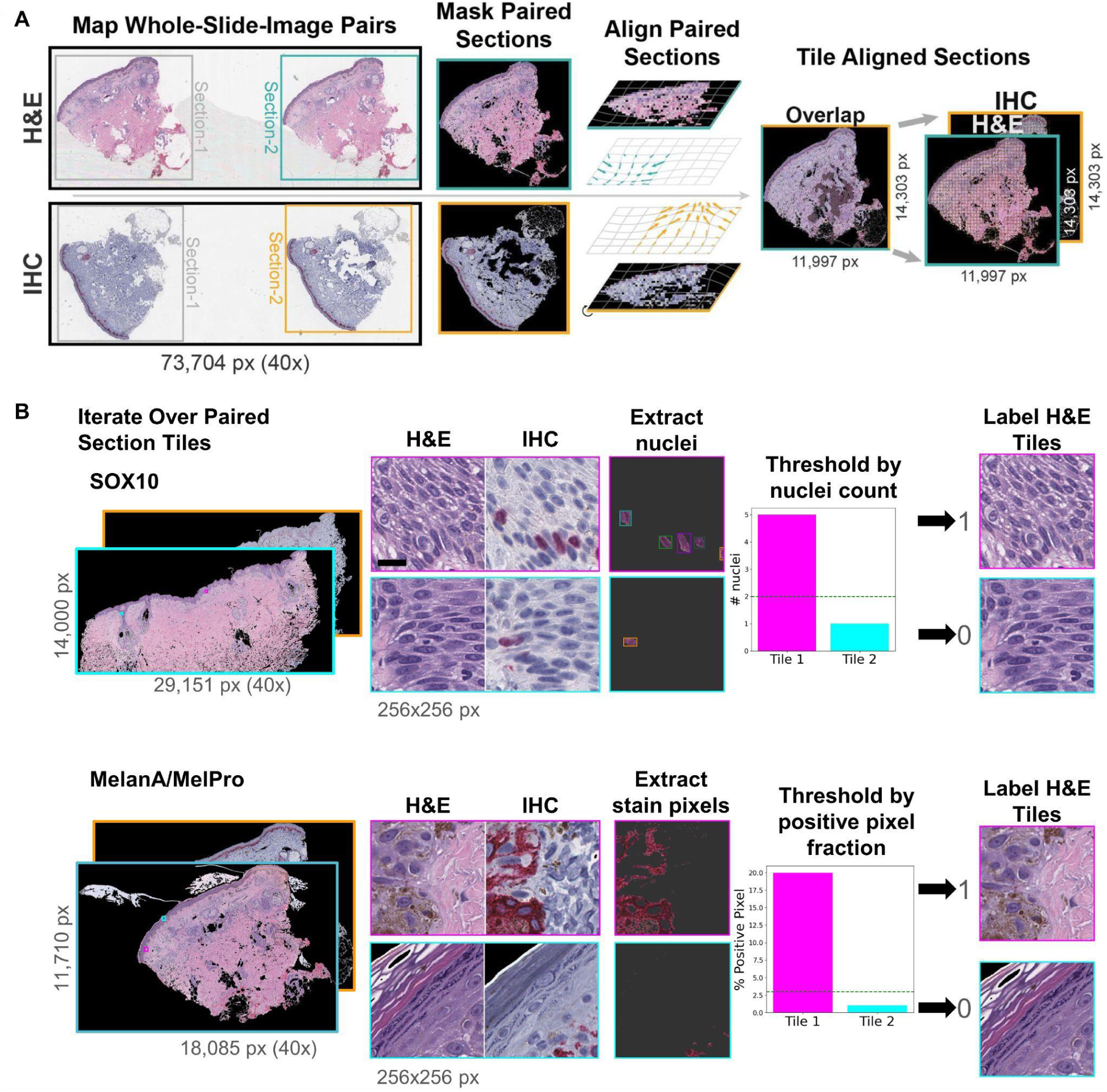
Whole-slide image processing pipeline: A) Tissue Section Preparation and Alignment: We extract and mask matching tissue sections from separate H&E and IHC-stained WSIs, forming a pair. We align tissue sections using Image-J’s b-spline alignment algorithm (BunwarpJ) to ensure optimal congruence at the tile level. Following alignment, we independently tile tissue sections. B) Tissue Tile Labeling: We label H&E tiles based on the associated IHC stain. For tiles corresponding to SOX10 stain, labeling is based on the count of nuclei, while for those with MelanA stain, the proportion of positive pixels. Scale bar = 16 μm.

We assess the performance of the (H&E) trained networks by their ability to classify tiled images from validation H&E patient tissue sections as either melanocyte-containing or non-melanocyte-containing, using unseen aligned IHC-stained tissue as ground truth. We generate high-resolution prediction heatmaps on validation tissue sections (H&E) for slide-level interpretation. We also perform an AI/ML explainability technique called saliency mapping to generate fine-grained maps highlighting which tissue regions contribute the most to the neural network’s detection of melanocytic tissue.

Teaching a supervised model to learn where melanocytes are present in an H&E tissue section, with confirmation solely from approximate information from a corresponding IHC stain, helps to reduce the necessity, cost, and time of manual dermatopathologist labeling and avoids the issues of human bias and noisy labeling^25,26^. We hope this “virtual stain” method will assist the general clinical pathologist in rapidly and effectively identifying cutaneous melanocytes, improving turnaround times, and reducing healthcare costs for new cutaneous MIS diagnoses.

## Methods

### Sample cohort

We generated digital pathology images from archival patient H&E and IHC stained slides of diagnosed melanoma in situ from either UCSF’s Dermatopathology Service hospital cases or Stanford’s Dermatopathology Service hospital cases. All samples were gathered between 2011-2015, with patients aged between 43-73. Each patient sample comprised a pair of WSIs: a WSI of H&E-stained tissue sections and a WSI of corresponding IHC-stained sections. At the original diagnosis, pathologists extracted and digitally scanned all available slides from various biopsies, including/incisional, shave, and punch biopsies. However, only H&E and IHC WSI biopsy pairs optimized for nearness based on alignment were included in the study.

### Sample preparation

Each cutaneous biopsy or excisional tissue underwent standard processing for H&E and IHC stained sections at time of original histopathologic diagnosis. For H&E staining, patient samples were formalin-fixed, sectioned, paraffin-embedded, cut and placed on a glass slide, and stained with hematoxylin and eosin (H&E). For IHC staining, tissue samples from the same paraffin blocks were similarly prepared but treated with one of three antibodies to highlight melanocytes: MelanA, SOX10, and MelPro (dual MelanA and Ki-67 stain). Some pairs were cut in exact serial tissue sections, and others were cut on a separate day after the tissue block was refaced at the discretion of the original diagnostic pathologist. We scanned each clinical case’s original diagnostic H&E and IHC glass slides to generate the WSI pairs. We digitized whole slide images at 40x magnification, corresponding to a resolution of 0.25 microns per pixel (MPP), using a Philips slide scanner (Philips Ultra Fast Scanner Research Use Only/ UFS1.6 RUO) or an Aperio scanscope (AxioVision; Leica Biosystems) and stored them as compressed pyramidal image files (BigTIFF and SVS images).

### Dataset curation

Of an original dataset of 242 newly scanned cases of melanoma, we selected 61 WSI pairs (122 WSIs) from the UCSF and Stanford clinical archives. Each pair comprises a WSI containing an H&E-stained slide and an independent but complementary WSI IHC-stained slide. Of 61 candidate pairs, we removed 38 pairs by manual inspection due to poor data quality, such as folded, damaged, or blurry tissue or irreconcilable size differences between H&E and IHC stained tissue sections. The final dataset consisted of 23 patient WSI pairs (46 WSIs), including 10 SOX10 stains, 10 MelanA stains (also known as MART1), and 3 MelPro stains (Supplemental Figure 1, Supplemental Tables 1, 2, and 3 detail dataset composition). We created two datasets by grouping patient pairs across institutions (UCSF: 14/23 samples; Stanford: 9/23 samples) by their IHC stains: one comprising WSIs with either MelanA or MelPro IHCs (combined: 13/23 WSIs), named the “MelanA” dataset, and a “SOX10” dataset (10/23 WSIs). SOX10 is a nuclear immunohistochemical stain, MelanA is a cytoplasmic immunohistochemical stain, and MelPro is a double-stain that highlights mitotically active nuclei (Ki67 stain) within the cytoplasmic-stained (MelanA stain) melanocyte population.

For morphologically diverse negative training examples, we randomly collected 45 non-cutaneous H&E tissue samples from a subset of TCGA to create 135,000 negative labeled tiles^32^ (Supplemental Table 4). These negative controls focused on pancreatic, breast, and lung carcinomas (Supplemental Table 5). The rationale for adding diverse negative H&E examples was to improve model generalization and ensure it does not simply learn to over-interpret hematoxylin as a surrogate for melanocytes.

### Tissue extraction and filtering

To automate tissue extraction, we developed a custom toolkit in Python (Supplemental Figure 2). The toolkit first identified the HSV color space of a WSI’s background. Using this information, it selected candidate tissue foreground regions that met a size requirement.

Thresholds for chroma, size, and degree of region-enclosing were manually adjustable, allowing for optimal tissue extraction. After the user refined the H&E and IHC WSI tissue regions, the toolkit matched tissue section pairs. Matching occurred automatically based on tissue locations in the WSI and could be manually adjusted by the user if necessary. Cases where the matched H&E or IHC section had digital artifacts, excessive tissue damage, was missing, was too small, or was indistinguishable from the background were filtered out. The toolkit then individually cropped and masked the background of successfully extracted tissue sections. The toolkit employed the open-source libraries pyvips^33^, OpenCV^34^, and IpyWidgets^35^.

### Image alignment

We used archival IHC-stained tissue sections as proxy melanocyte labels for matched H&E sections. This method required that adjacent sections were morphologically similar and that a proxy label from one section relayed information about the other. Further, at the local level, the density of IHC staining for melanocytes was correlated with the probability of melanocytes being present within the same region of the adjacent H&E slice. As models operated on high-magnification H&E inputs at the 256x256 pixel level, alignment between H&E and IHC tissue sections was crucial.

We align matched H&E and IHC tissue sections at the section-pair level, at native resolution (40x magnification; 0.25 MPP), using the open-source alignment algorithm bUnwarpJ^141^ (Figure 1a). In brief, as calculated by an energy function, the algorithm registered image pairs using 2D elastic deformations (B-splines) to minimize differences. Matching tissue sections often varied in shape and size after extraction from WSIs. The algorithm zero-padded (i.e., fills in gaps with black pixels) the smallest section from each matched pair to the size of the larger section before alignment. Additionally, we cropped sections otherwise too large to align (>5GB) to create two smaller pairs instead of one large pair. We manually inspected alignments, scored them, and considered coefficients returned from the alignment algorithm. Alignment coefficients and manual inspection scores were similar (Supplemental Tables 2 and 3). We discarded candidate pairs that were unalignable or received a low alignment score.

### Stain-specific labeling

We used multiple IHC stains that operated as different melanocytic biomarkers: nuclear and cytoplasmic. We used red and brown chromogen SOX10 and red chromogen MelanA and MelPro stains. Using computational color deconvolution, we converted red-green-blue (RGB) IHC-stained images into an intermediary hematoxylin-eosin-diaminobenzidine (HED) color space. The brown color of 3,3’-diaminobenzidine (DAB) denoted antibody expression for MelanA, SOX10, or MelPro.

Before isolating the DAB, we removed major artifacts, such as blue or green ink and tissue areas containing hemorrhage (Supplemental Figure 3, ink example). Otherwise, artifacts exhibited strong “false positive” DAB signal indistinguishable from true melanocyte expression. We applied a sequence of Otsu thresholding steps to segment candidate artifact regions^36^ and removed ink features using predefined hue-saturation-value (HSV) color exclusion ranges. Furthermore, we manually corrected mislabeled non-melanocyte regions identified as melanocytes (Supplemental Figure 4).

Following artifact removal, we separated DAB signal from the IHC stain images to extract nuclei for SOX10 and cytoplasmic pixels for MelanA stains. We kept the resulting images at 40x magnification. We generated a training dataset for the models by partitioning the images into 256x256 pixel tiles, with a stride (overlap) of 128 pixels for training and 256 pixels for validation and test data.

We used positive-signal threshold calculations specific to the melanocytic biomarker type (Figure 1b; see GitHub open code repository). For SOX10, we labeled H&E tiles positive if their matched IHC tile contained more than two discrete DAB-positive nuclei. For MelanA, we required that DAB-positive pixels comprised more than three percent of the tile area. All H&E tiles failing these criteria received a melanocyte-negative training label. We established thresholds by a board-certified dermatopathologist (ESK) manually inspecting the tiles and their labels across a range of potential values (Supplemental Figure 5). In an inherent biological limitation, despite their widespread diagnostic use, none of the IHC stains are exclusive to melanocytic cells. For instance, SOX10 also stains dermal nerve bundles and eccrine glands, but it is considered specific for melanocytes within the epidermis.

### Convolutional neural network model training and assessment

As the archival slides originally served clinical diagnostic needs, generating high-magnification and relevant paired WSI training data post hoc often raised challenges. We consequently had to discard a substantial portion of the dataset during preprocessing, necessitating operating in a low-data regime for deep learning. Consequently, we adopted a training and evaluation strategy motivated by 5-fold cross-validation, with splits into 60% training, 20% validation, and 20% testing data (tiles, stratified by patient case) for each fold.

We injected TCGA negative tiles into training (70%) and validation (30%) batches, using a batch size of 240 tiles (Supplemental Table 6). We normalized H&E image pixel input ranges by subtracting their mean value and dividing by the standard deviation of the training set. We trained independent stain-specific convolutional neural networks (CNNs) using a DenseNet121 architecture with early stopping using a patience of 8 epochs. We used a stochastic gradient descent (SGD) optimizer with a learning rate of 1e-4 after a grid search hyperparameter optimization that explored combinations of optimizer = {SGD, Adam} and learning rate = {1e-5, 3e-5, 1e-4, 3e-4, 1e-3, 3e-3}. We assessed performance by the CNNs’ ability to predict H&E tile labels derived from their unseen IHC counterparts (see “Stain-specific labeling” above). We measured the area under the receiver operating characteristic (AUROC) and the area under the precision-recall curve (AURPC) for each model on its representative test hold-out set per fold and reported the all-fold averages as SOX10 and MelanA CNN performance.

### Prediction heatmaps and IHC label maps

We generated prediction heatmaps to visually summarize the CNN’s confidence of melanocyte presence across entire H&E tissue sections using a custom multiprocessing script (see GitHub repository) with PyTorch^142^. We generated 256x256 tiles centered on every 6^th^ pixel (i.e., stride=6). We zero-padded tiles to reach the appropriate size, when needed, at the edges of the image. The CNN calculates a prediction score on each tile independently. We converted and normalized prediction scores to RGB values in the matplotlib “viridis” color space and visualized them over the original H&E image using Python.

By contrast, IHC label maps illustrate the “best case” visualization of melanocytic-positive labels on tissue sections, accounting for the visual smoothing that inherently arises from 256x256 pixel (0.25 MPP) tiles when displayed at the same 6-pixel stride. These maps derive directly from the hidden DAB-based labels (see “Stain-specific labeling” above) and represent what a “perfect” prediction heatmap could achieve. Generating label maps follows a similar process to prediction heatmaps, except that instead of using the trained CNN to get a prediction based on the H&E, the labeling method highlights the tile-smoothed actual DAB-positive melanocyte area (yellow overlay) on the IHC images.

### Generating saliency and agreement maps

Saliency mapping methods highlight the cytologic and architectural pixel features a CNN finds important within an H&E input image. To examine which pixels in an H&E tissue section achieve the highest attribution values numerically, we used the Captum library^37^ to apply guided gradient-weighted class activation mapping (Guided Grad-CAM^38^) and integrated gradients^39,40^ on all convolutional layers of DensetNet121^31^. We also used the Captum library’s “NoiseTunnel” to generate five samples of the input tile with Gaussian noise to allow for more stable attributions. We used Guided Grad-CAM’s attributions in Figures 5 and 6 because they were more visually distinct than those from integrated gradients (Supplemental Figure 7).

After generating attributions for each convolutional layer of the model, we averaged these and overlaid them on the H&E images using the “viridis” colormap (Figure 6a) and custom versions of the reversed “plasma” colormap for the “explained-heatmap” overlays (Figure 6b; Figure 6c, right). The “agreement map” (Figure 6c, left) used a custom colormap with solid colors assigned to the Agreement, MelanA, and SOX10 categories. Figures 6b and 6c’s H&E color signal is also conditioned on the models’ positive predictive signal (model confidence threshold=0.9). In other words, grayed areas are not predicted to have MelanA or Sox10 signal, per their original matched stain. The “explained-heatmap” and “agreement maps” are stitched views 1024x1024px or 4x4 tiles in size. Because attribution values can vary widely between tiles for differing models, we employed Captum’s attribute normalization feature for consistent comparison. We also applied Gaussian smoothing to remove edge effects between tiles. We used Python matplotlib, numpy, and OpenCV libraries to generate the images.

## Results

MelanA and SOX10 IHC stains are among the most common in melanocytic dermatopathology. We created two separate convolutional neural networks (CNN), trained each on a specific stain, and evaluated how they agreed and differed.

### We algorithmically generated stain labels from IHC sections aligned to H&E sections

We developed a stain-agnostic melanocyte presence labeling pipeline that annotated whether a small (64x64 micron; 256x256 pixel) H&E tile contained melanocytic cells using sequentially sliced matched nuclear and cytoplasmic IHC stained tissue images. Using adjacent IHC stained tissue as an approximate answer or “label” complements studies that rely on collecting training data from pathologists’ manual annotations. Despite the physical differences between different tissue sections, we demonstrated a computationally efficient method to label melanocytic cells in archival H&E and IHC WSIs accurately. Because our dataset is archival, the H&E sections were not always “back to back” or cut strictly adjacent to the IHC sections and may vary slightly in architecture and morphology. Consequently, a raw IHC WSI file, spanning tens of thousands of pixels in height and width, could not serve as a proxy to label for melanocytic cell presence to a matched archival H&E without further image processing.

Accordingly, we first separated the WSI into individual tissue sections to remove slide white space and to account for variations in tissue section placement or orientation (see Tissue Extraction and Filtering, Methods). Next, we tackled coarse-grained H&E-to-IHC section dissimilarity by algorithmically aligning the position, orientation, and scale of the IHC section to match a paired H&E section using publicly available software, bUnwarpJ (Figure 1a). This tool also computes limited “warping” corrections, such as adjusting for minor shearing or bunching of the tissue samples. High-magnification differences typically persisted between H&E and IHC tissue sections, as evident when comparing alignment fidelity within 256x256 pixel H&E and IHC tiles (Figure 1b). However, the exact variation in tissue structures across tiles became insignificant when we consolidated the IHC tile’s signal into a single binary annotation per H&E tile (Figure 1b, rightmost columns). Consequently, the neural networks never received within-tile label information during training. However, this inherent limitation to the slide processing methods did restrict our pipeline to processing WSIs at a maximum optical magnification of 40x (0.25 microns per pixel, MPP), as the alignment limitations did not support a finer-grained training label.

We developed separate methods to label H&E tiles by paired SOX10 stain tiles versus MelanA and MelPro stained tiles, tailored to the IHC stain’s biological characteristics and to facilitate downstream *in silico* multiplex melanocytic staining. SOX10 is a nuclear stain, so we counted stained nuclei. Since MelanA and MelPro are cytoplasmic stains, we followed a more cytoplasmic logic for these stains (hereafter simplified to “MelanA”) by calculating the proportion of stain-positive pixels within an IHC tile. We set positive-label thresholds by dermatopathologist review (ESK), requiring at least two stained nuclei for SOX10 and three percent of the pixel area to be stain-positive for MelanA (Figure 1b). As expected, increasing these thresholds reduces the total count of positively labeled tiles while lowering them has the opposite effect (Supplemental Figure 5).

### Models detected melanocyte cells in H&E-stained tissue

We created *in silico* multiplex IHC stains from archival H&E tissue by developing convolutional neural network (CNN) models that assessed the likelihood of melanocytic cell presence one H&E tile at a time. We trained DenseNet121 CNN models^31^ on H&E tiled images labeled from the MelanA and SOX10 datasets. Many of the archival WSIs we initially collected failed in the data processing step due to insurmountable tissue-structure differences between H&E vs IHC sections or digitization artifacts such as blurriness – we discarded 62% of the original 61 H&E-IHC WSI pairs (Supplemental Table 2 and 3 “Action” column). Consequently, a case-stratified static hold-out test set strategy was not feasible at this dataset size. To evaluate the models reliably, we performed 5-fold cross-validation and reported the average performance among the folds.

MelanA and SOX10 models achieved strong AUROCs (0.948±0.023 and 0.867±0.091) with good AUPRCs (0.611±0.091 and 0.433±0.165) (Figure 2a). We also constructed “label maps” directly on the hidden IHC tissue sections to visually evaluate this numerical performance. The logic was that the training pipeline, by definition, could not generate cell-level melanocyte predictions because we could only train the CNNs to generate binary yes/no predictions at the level of 256x256 pixel (∼ 0.5x0.5 micron) fields of view. While we could progressively “slide” the CNN’s field of view one pixel at a time across the entire H&E section, we still needed to average the resulting overlapping predictions to generate the “prediction heatmap” (Figure 2b, middle column). Consequently, even a “perfect” CNN could not generate heatmaps more fine-grained than the analogous overlapping sliding-window procedure on the hidden paired IHC, which we termed “label maps” (Figure 2b, right column). Whole-tissue prediction heatmaps achieved excellent fidelity to the IHC label maps, particularly in the high-confidence (dark purple overlay) regions (top row, Figure 2b and Figure 3b). In the zoomed insets (bottom row, Figure 2b and Figure 3b), both models appear slightly oversensitive in their melanocytic cell area prediction, consistent with the tradeoffs shown in their precision-recall curves (right, Figure 2a and Figure 3a). Naive 1-pixel-at-a-time (“1-pixel stride”) heatmaps require 10,000^2^ = 100 million CNN predictions per 10k-by-10k pixel WSI, so we generated whole-tissue heatmaps using a 6-pixel stride and reported dataset-wide performance metrics using a 256-pixel stride. Performance correlated closely between 6-pixel and 256-pixel strides (Supplemental Figure 6).

**Figure 2.**
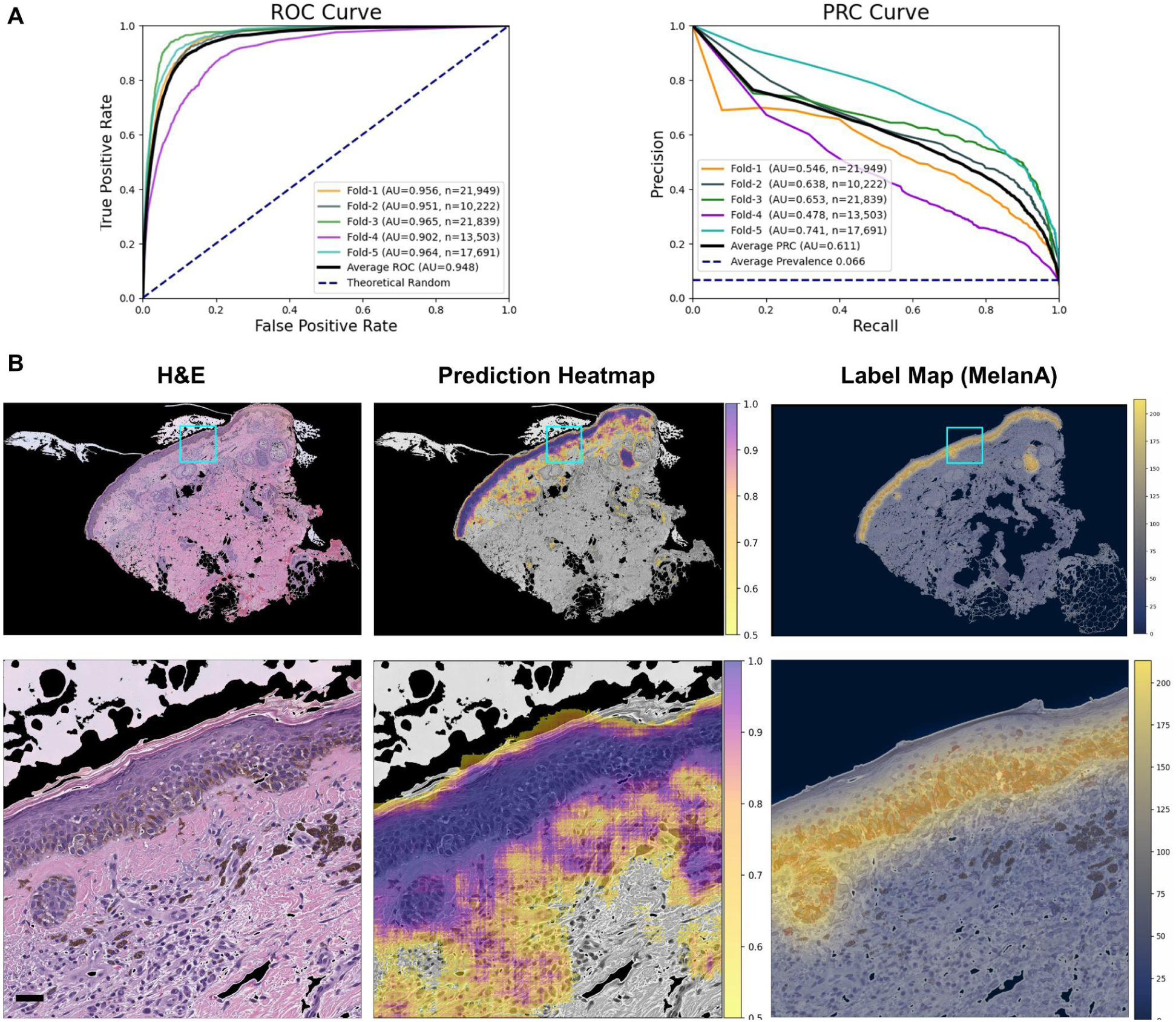
MelanA melanocyte-detector evaluation. A) Performance Metrics: We evaluate the models, labeled based on either MelanA or MelPro IHC stains, using five distinct test sets from a 5-fold cross-validation. Each fold splits patients into train vs testing categories. The AUROC (left) and AUPRC (right) curves display the performances across these test sets. B) Visualizing Melanocytes Predictions: The top row compares the original H&E image, the prediction heatmap, and the corresponding ground-truth IHC label map. The bottom row shows a zoomed-in, higher-resolution segment of the top row image, highlighted in blue. Scale bar = 32 μm.

**Figure 3.**
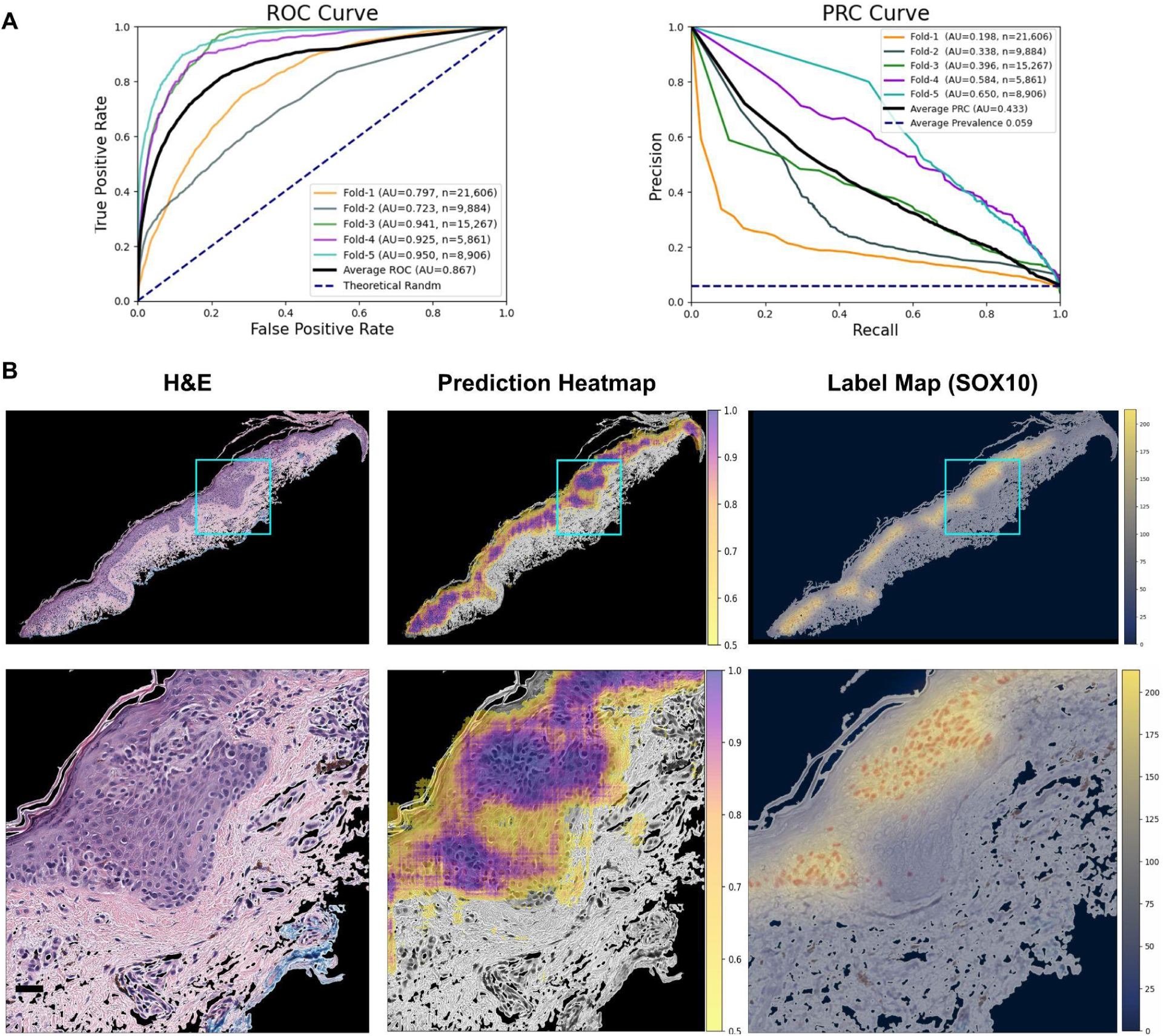
SOX10 melanocyte-detector evaluation. We prepared panels (a) and (b) as in Figure 2, but using SOX10 IHC stained whole slide images. Scale bar = 32 μm.

### Models accurately identified melanocytes, with biologically sensible edge cases

We used validation tissue tiles of H&E stained sections paired with IHC stained sections to test the MelanA and SOX10 models. Overall, the models accurately and reliably identified H&E-stained tiles with increased numbers of melanocytes (Figure 4).

**Figure 4.**
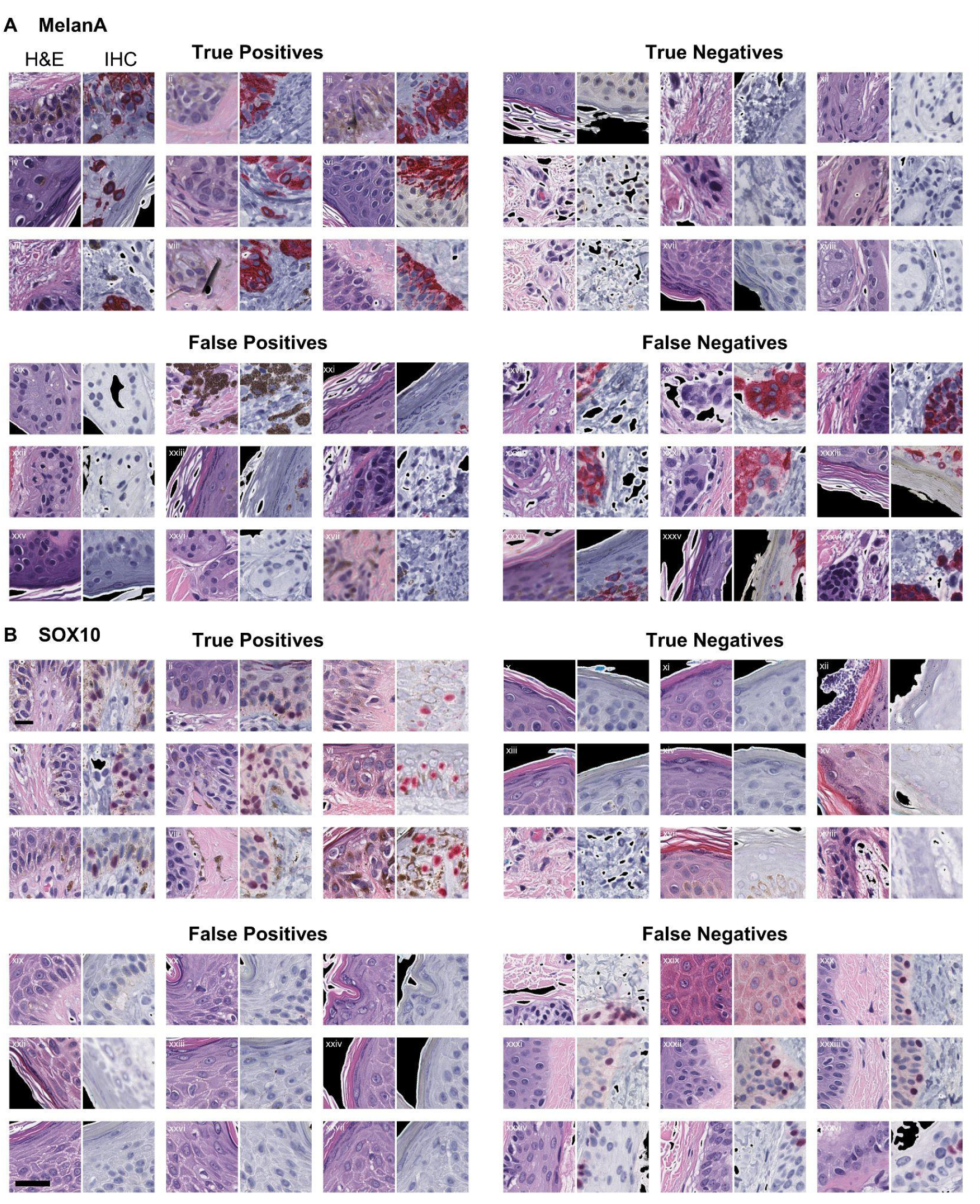
Assessing model accuracy and reliability in identifying melanocytes in validation H&E and IHC stained tiles. Truth Table Prediction Examples: A and B) Predictions of MelanA and SOX10 Melanocyte-Detectors: Both panels demonstrate the four prediction outcomes: true positive, true negative, false positive, and false negative. For each outcome category, nine paired examples contain a paired H&E tile and its corresponding IHC tile for the MelanA (A) and SOX10 (B) models. Scale bar = 32 μm.

Melanocytes show a spectrum of morphologies, from small hyperchromatic nuclei with perinuclear clearing to enlarged nuclei with nucleoli and characteristic “battleship gray” cytoplasm^41^. Some melanocyte cytologic features overlap with keratinocyte cytologic features, making cell-specific identification particularly challenging in H&E stained images. We wondered whether the SOX10 and MelanA CNN models had learned to leverage these characteristic morphological features or instead relied on an unrelated numerical logic. To assess this, we examined prediction successes and failures at the single-tile level (256x256 pixel tile with 0.25 MPP, so 64x64 microns, Figure 4). We display a representative subset of true positive, true negative, false positive, and false negative tiles for each model, arising from tile-level predictions of whether predicted melanocyte counts exceeded a minimum threshold. In the previous section, the prediction heatmaps (Figure 2b and Figure 3b, top row) visualize averaged prediction confidences by overlapping individual predictions for these high-power (40x) but small field-of-view (64x64 micron) tiles to simulate a low magnification view of the entire tissue. MelanA and SOX10 models accurately identified tiles with increased numbers of melanocytes, including tiles with subtle melanocytes interspersed within the epidermal keratinocytes (e.g., “True Positives” in Figure 4a,b).

Differentiating eccrine glands from melanocytic nests was more challenging for the models than it typically would be for pathologists. Eccrine glands display small hyperchromatic nuclei and amphophilic cytoplasm, similar to some melanocytes, but their duct and gland architecture plus deep dermal location are generally clear non-melanocytic indicators to pathologists. SOX10 is known to stain eccrine myoepithelial and secretory cells, whereas MelanA does not. The MelanA model incorrectly identified some eccrine glands as melanocytes (e.g., “False Positives” tiles *xix, xxii, xxvi* of Figure 4a) but still appropriately identified other “true negative” eccrine gland tiles (e.g., *xii, xv, xviii* of Figure 4a). Since the SOX10 model inherits the limitations of the SOX10 antibody by its training, it is unsurprising that this model also identified some “true positive” tiles that contain eccrine glands instead of melanocytes.

Consequently, numerical performance scores for the SOX10 model (e.g., Figure 3a) formally reflect prediction of positive IHC signal instead of melanocyte-specific signal. However, the model explainability methods (Figure 5) reveal that both models nonetheless assign similar importance to nuclear and cytoplasmic features in eccrine glands in H&E stained images despite their positive SOX10 and negative MelanA expression (“False Positives” tile sets *iv* and *vi* in Figure 5b and “True Negatives” tile sets *ii* and *iii* in Figure 5c).

**Figure 5.**
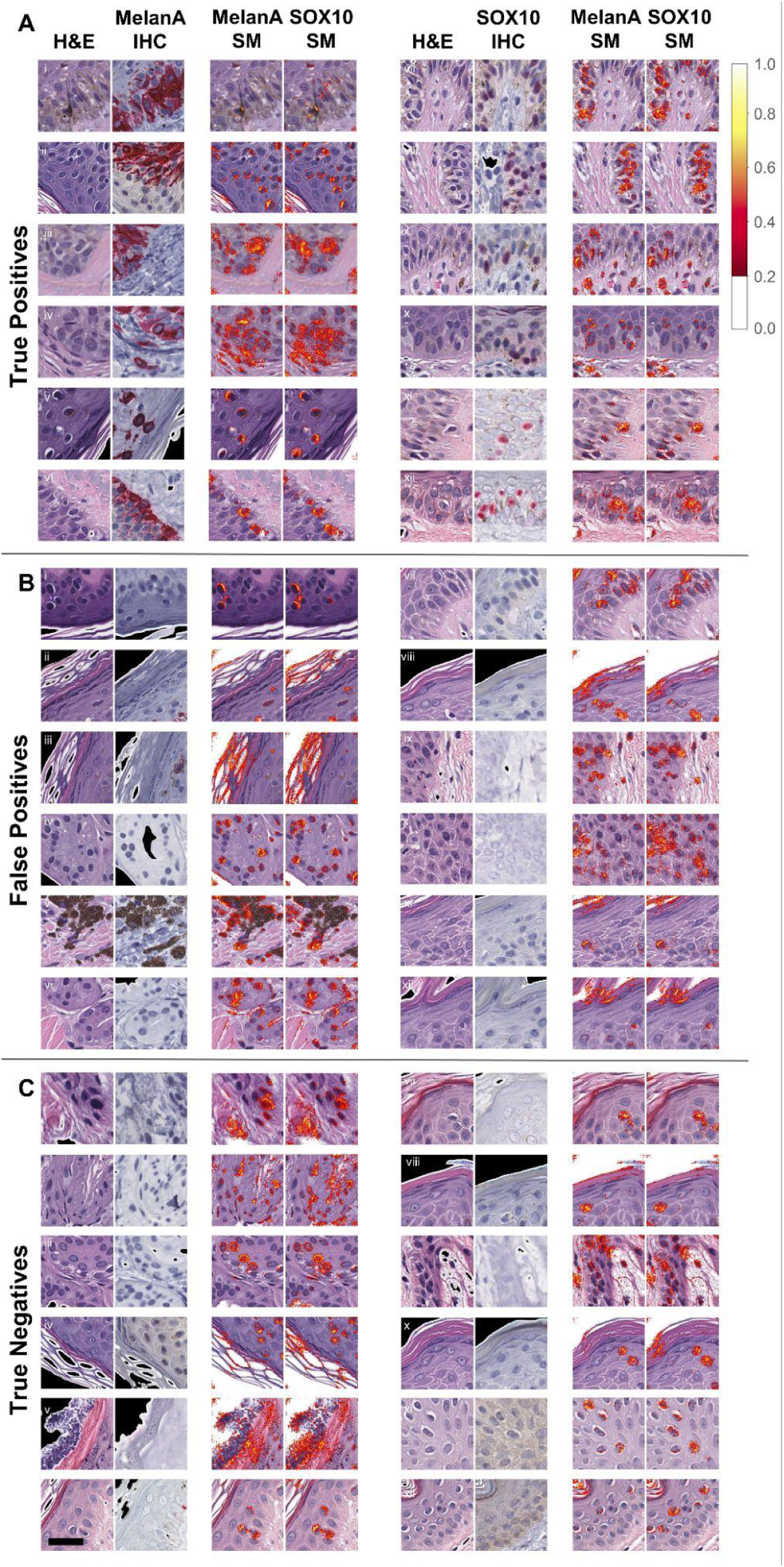
Example of saliency map on true positive (A), false negative tiles (B), and true negative example tiles (C) using the MelanA and SOX10 models. Scale bar = 32 μm.

The SOX10 model incorrectly rejected some tiles with at least three melanocytes in the H&E image despite the image pairs being back to back, with the same cells in both images. In these scenarios, the algorithm did not correctly identify the small hyperchromatic melanocytes (e.g., “False Negatives” tile *xxx*, Figure 4b). However, other “false” negative image pairs appear to be cut discontinuously, with either fewer or absent melanocytes seen in the H&E half of the tile pair or a blurry H&E image (“False Negatives” tile *xxxi, xxxii, xxxiv, xxxvi* Figure 4b). These discontinuous cases inflate the “false negatives” count, as the model reliably evaluated the H&E tile. In contrast, the MelanA model, on average, rejects tiles with more melanocytes than the SOX10 tiles, but there appears to be more processing artifact within the tissue in these images and non-contiguous section “mismatch” in some cross-tissue tile alignment (e.g., “False Negatives” tiles *xxxi, xxxvi* in Figure 4a). Therefore, some “false negative” MelanA examples may be misleading.

MelanA and SOX10 models both appeared to select tiles with basketweave orthokeratosis and few to no melanocytes in their “false positives” (e.g., “False Positives” tiles *xxi, xxiii* in Figure 4a; “False Positives” tiles *xxii, xxiv* in Figure 4b). Some melanocytic lesions may show pigmented parakeratosis, such as in acral sites or particularly irritated or “traumatized” melanocytic lesions, but basketweave or compact orthokeratosis is not conventionally independently correlated with the presence or absence of melanocytes. Further, the stratum corneum in false positive tiles without melanocytes appears to lack obvious pigmentation. While the basketweave orthokeratosis could be coincidentally present in these “false positive” tiles, its saliency highlighted in Figure 5 argues against mere chance (“False Positives” tiles *ii, iii, viii, xi, xii* in Figure 5b).

Cases where the SOX10 model falsely identified keratinocytes as melanocytes show H&E tiles with perinuclear clearing of keratinocytes, particularly in cases with angulated and hyperchromatic keratinocytic nuclei (e.g., “False Positives” tiles *xx, xxvi* in Figure 4b). These cases also show epidermal “spongiosis,” signifying increased intercellular fluid, creating visible desmosomes between cells and “reactive” keratinocytes. The selection of tiles with keratinocyte hemi-desmosomes is a surprising finding, given that melanocytes lack desmosomes. However, these reactive keratinocytes contain more prominent nucleoli than their neighbors, and some cells show nuclear pallor mimicking melanocyte pseudo-inclusions, suggesting the SOX10 model may be prioritizing nuclear features and immediate perinuclear features of the cytoplasm rather than the intercellular bridging seen elsewhere in the image (see saliency in “False Positives” tiles *vii, ix, x* in Figure 5b).

### Interpreting the models’ rationales for melanocytic morphology

Intuitively, a saliency map illustrates a model’s rationale for predicting melanocyte presence or absence in the tiles. Specifically, the maps highlight a tile’s most “salient” features (pixels), whose deletion or intensification would change the model’s prediction most. For the clinically minded, these salient features are pertinent positives *and* negatives, with the saliency maps highlighting features in the image that can also explain why something *isn’t* a melanocyte. Unlike the heat maps showing the probability of increased melanocytes in tiles (e.g., Figure 2b and Figure 3b), the saliency maps highlight the specific pixels the algorithm uses to decide whether or not increased numbers of melanocytes are present. In some cases, the algorithm identifies features inside a melanocyte as contributory. In other cases, the model identifies important discriminatory features elsewhere in the image, such as within keratinocytes or the stratum corneum.

Saliency mapping techniques calculate a numerical contribution strength, or “attribution,” for each pixel (feature) within a model’s prediction *context*. Thus, a salient pixel will have a high attribution value by a particular model in the context of a particular prediction. The same pixel may lose salience in the context of a different model (e.g., a model trained on a different IHC stain) or a prediction on an overlapping but different tile. We used Guided Grad-CAM (see Methods) to calculate and visualize attributions on the H&E input tile using SOX10 and MelanA models for true positive and false positive outcomes (Figure 5; Saliency Maps, SM). We scaled attribution magnitudes between 0 and 1, with attribution strength proceeding from red (weak, but >0.2) to yellow (strong). For comparison, Figure 6a displays the raw output of Guided Grad-CAM alongside its H&E instead of by overlay (rightmost two columns, titled “SM”). We generated independent saliency maps for the MelanA- and SOX10-trained models (Figure 5, rightmost two columns of each image group). Despite meaningful differences in how we calculated H&E tile labels when training each model, the saliency maps nonetheless appear surprisingly similar. The SOX10 model relied on a count of stained positive nuclei within the hidden paired IHC, whereas the MelanA model learned from labels based on the proportion of stained positive cytoplasmic areas. Both models learned to assign attention to similar cell morphologies of the H&E-stained input tissue.

**Figure 6.**
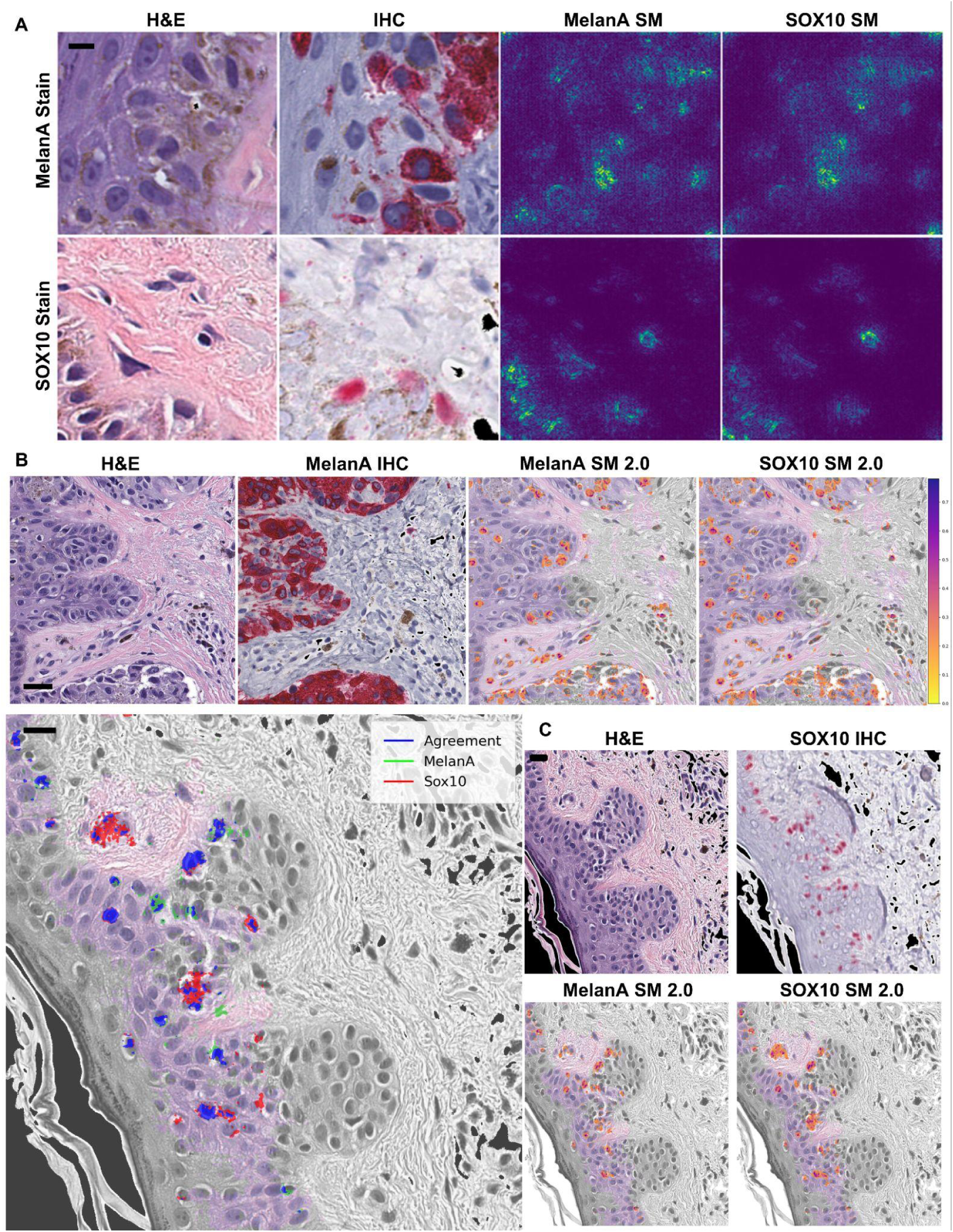
Interpreting model predictions via saliency mapping. A) Columns display H&E tiles, IHC, MelanA model saliency map attributions, and SOX10 model saliency map attributions. All tiles are size 256x256 pixels, scale bar = 8 μm. B) A stitched view of attributions across multiple tiles (1024x1024px) on a MelanA-stained tissue. We used Guided Grad-CAM to calculate attributions and overlaid them onto the portion of the tissue in the image predicted to have melanocytes (the color-saturated area; using prediction heatmap, stride=6, confidence threshold=0.9). The desaturated area represents tissue without strong melanocyte prediction. Scale bar = 32 μm. C) We constructed a multi-antibody “agreement map” view to compare the virtual stains (left), indicating how the rationales of the two models compare. Both models independently assign high attribution to blue pixels, whereas MelanA-model-only attributions are green, and SOX10-model-only attributions are green. We include corresponding H&E, IHC (top right), and separated saliency maps (bottom right) for reference. The desaturated regions again correspond to the region lacking strong SOX10 prediction (using prediction heatmap, stride=6, confidence threshold=0.9). Scale bar = 16 μm.

The MelanA model identifies portions of the stratum corneum and dense dermal melanin deposition as salient features, albeit incorrectly in some cases (e.g., tile sets *ii, iii, and v* in Figure 5b). However, it correctly disregards increased intracytoplasmic melanin pigment in other less densely pigmented cases, instead focusing on the nuclear membrane and perinuclear salient features (e.g., tile sets *vii-xii* in Figure 5a). In instances where the models correctly identify a tile as lacking increased melanocytes, it identifies bizarrely enlarged nuclei and correctly identifies keratinocytic intracytoplasmic processes (desmosomes) as specifically contributing to its correct tile interpretation (e.g., tile sets *vii* and *viii* in Figure 5c). Interestingly, both models appropriately identify nuclear membrane and perinuclear features within eccrine glands as salient reasons for the tile *not* containing increased numbers of melanocytes (e.g., tile sets *ii* and *iii* in Figure 5c).

### Nuclear and perinuclear features are identifying features of melanocytes in both models

Prediction heatmaps (e.g., Figures 2b and 3b) show areas of increased melanocytes, while saliency maps (e.g., Figures 5 and 6a) show the most essential pixels underlying the predictions. We posited that combining these maps into a single view would help researchers and clinicians visualize neural network predictions without requiring that they accept all model predictions in a “black box” way. To achieve this, we generated an “explained-heatmap” that highlights increased-melanocyte regions (colored areas vs. grayscale) and calls out the pixels motivating the positive prediction for those regions (orange) (Figure 6b). By conditioning the saliency map (yellow-orange) on positive prediction-only regions, the explained-heatmap focuses on areas and visual morphology reasoning where the model predicts positive outcomes with above 0.9 confidence. Intriguingly, this pixel-level reasoning extracted from the saliency mapping techniques is a byproduct of the neural network training process, which never received training labels at a resolution more fine-grained than yes or no labels for entire 256x256 pixel input images of H&E tiles.

On inspection, explained-heatmap views for MelanA- and SOX10-based models appeared unexpectedly similar (e.g., Figure 6a and Figure 5). To distinguish subtle differences between the models trained on labels from these different IHC stains, we also generated a virtual-IHC “agreement map” comparison view (Figure 6b). Green and red overlays call out pixels exclusively salient to MelanA or SOX10 model predictions on the H&E image. Where both models agreed a pixel was salient, the color is blue. Although MelanA is a cytoplasmic stain and SOX10 is a nuclear stain, the models’ attributions “converge” on salient features of the nucleus and the nuclear-cytoplasmic interface, as well as immediate perinuclear cytoplasmic features. Although stratum corneum and intercellular features were salient in negative predictions (e.g., Figure 5c), neither category appears to predict the presence of melanocytes in these positive-area examples (Figure 6c).

## Discussion

Convolutional neural networks (CNNs) trained on a small real-world dataset of approximate and noisy H&E- and IHC-stained archival slide pairs nonetheless learned robust and consistent visual reasoning (pixel salience) for melanocyte cell presence. Although the CNN models never received training information more granular than a simple “melanocytic” or “not” label for 64x64 micron H&E tiles, they learned to ascribe precise and fine-grained salience to specific cells (Figure 5). In evaluating the CNNs’ reasoning for melanocyte-positive area predictions, we found the salient cells were morphologically plausible (Figure 5a, Figure 6a-b).

Strikingly, independent CNNs trained on different IHC stains nonetheless independently converged on similar visual reasoning (e.g., Figure 6c) despite the stains themselves being cytoplasmic (MelanA) vs nuclear (SOX10). This similarity may arise partly because the IHC-based training labels are coarse-grained (e.g., one binary label per 64x64 micron tile), but these labels are not interchangeable, and the models trained on them do not make identical predictions or exhibit identical reasoning. Consequently, the extent to which the IHC-specific models agree on a particular piece of tissue in predictions or calculated reasoning may also reveal a rudimentary measure of certainty.

Using adjacent but inherently different IHC-stained tissue to derive training labels tackles a problem at the heart of supervised learning – that outputs are only as good as their inputs. In pathology and computational models alike, the phrase “garbage in, garbage out” emphasizes poor quality data, such as from non-representative biopsy sampling, incomplete tissue preservation, or inconsistent labeling^42,43^, hampers diagnostic accuracy. However, the gold standard of expert human annotation to identify melanocytes remains unsatisfactory for precise computational labeling because humans can be inconsistent and lack perfect interobserver consensus even at the expert level^44–47^. These inconsistencies can arise from error, but also from differing areas of emphasis, expertise^48^, and positive-criteria definitions^49^. Consequently, we attempted to exploit biological multimodality as a surrogate for ground truth, accepting only the slide-wide binary human input that melanocytic atypia existed within a whole slide image. We then used immunohistochemical staining to label adjacent H&E sections to train a CNN of pathologist-diagnosed melanoma in situ images.

The resulting models learned features indicative of melanocyte presence (Figure 5, Figure 6) within H&E images and performed well on validation patient samples (Figure 2a, Figure 3a). Models are robust to similar-looking tissue areas that do not contain melanocytic cells (Figure 4) and can use information from different stains in alternative but complementary ways (Figure 6a). We can map our predictions back onto tissue samples to provide an interpretable view for pathologists to consider, and we observe what morphological features trained networks are activated by (Figure 5, Figure 6a-b). Despite overall small training dataset sizes and the biological differences between the different IHC targets MelanA (cytoplasmic) and SOX10 (nuclear), the independent CNN models learned remarkably similar pixel-wise rationales for their predictions (Figure 5, Figure 6c).

Saliency maps visually illustrate “why” a model predicts there are (or are not) increased numbers of melanocytes in an H&E input tile. For the clinically minded, these “salient features” can be thought of as both pertinent positives *and* pertinent negatives, with some of the features highlighted by the saliency maps representing specific features in the image that explain why an object *is not* a melanocyte. Similarly, pathologists develop a gestalt of when increased melanocytes are likely to be present in the epidermis based on the surrounding architecture, degree of basilar keratinocytic hyperpigmentation or pigmented parakeratosis, and presence or absence of a clinically-targeted keratinocytic neoplasm or inflammatory condition. The saliency maps are consistent with the computational models leveraging similar field-of-view level data when identifying features of the image that appear unrelated to a melanocyte, but this is only a hypothesis without further testing, such as synthetic counterfactual experiments.

Several caveats merit mention. Biologically, eccrine glands challenged both models. In future studies, additional training with a special stain, such as a Periodic acid-Schiff (PAS) stain^50^ highlighting secretory granules, may help the models differentiate between eccrine glands and melanocytes. Computationally, compared to some large-scale studies on tens to hundreds of thousands of private multi-tissue slides^15,51^, this melanocytic tissue dataset is small, varied, and visually noisy. It spans two disparate melanocytic IHC stains and medical institutions using different slide scanners and sensors. Indeed, the archival and real-world nature of the slide pairs meant that these biopsy cases were among the most pathologically complex, requiring the additional step of IHC slides for diagnosis. As we drew them from physical archives, the IHC slides were not always cut “back to back” and demonstrated a varying degree of fading due to time and storage conditions. We ultimately had to discard half of the digitized slides from model training due to image-quality concerns or the physical tissue sections being too different across the slides for meaningful comparison. Consequently, employing fine-grained deep learning models and strategies such as semantic segmentation (e.g., U-Net^52,53^) or object detection^54,55^ was impossible, including intriguing but data-hungry new architectures such as vision transformers^56^.

However, these apparent limitations became a strength of the study, as its most striking findings were the consistent and convergent nature of the two CNNs’ predictions and saliency maps despite the observation that deep learning models can fail to generalize in scientific and medical settings^57,58^. Some areas of diagnostic pathology, such as levels of melanocytic atypia, are not black and white – requiring technical skill and subjective expert opinion. Just as pathologists must make diagnostic decisions in imperfect real-world conditions, these computational models “learned” to grapple with imperfect inputs to provide reproducible tile-level melanocyte “labels” in various histopathologic digital images over time and across two different institutions. Future studies would ideally expand the models’ training with additional levels of atypia, akin to pathology residents learning more subtle disease variations after tackling overtly benign versus malignant examples.

## List of Abbreviations

ML: machine learning
DL: deep learning
WSI: whole slide image
GPU: graphics processing unit
MPP: microns per pixel
px: pixel
SOX10: SRY related HMG box 10 (SOX10) protein (IHC stain), nuclear stain
MelanA: Melanoma Antigen (IHC stain). Synonymous with MART1 (“Melanoma Antigen Recognized by T cells 1”), cytoplasmic stain
MelPro: MelanA plus Ki67-antibody MIB1 combination IHC stain, nuclear (Ki67) and cytoplasmic (MelanA)
CNN: convolutional neural network
H&E: hematoxylin and eosin
IHC: immunohistochemical
ROC: receiver operating characteristic
PRC: precision-recall curve
AU: area under
AUC: area under the curve
MIS: melanoma in situ
FFPE: formalin-fixed paraffin-embedded
SM: saliency map

## Declarations

### Ethics approval and consent to participate

We obtained all materials through pre-approved teaching slides from UCSF’s dermatopathology program or from patients who gave informed consent to distribute and use their samples at UCSF or Stanford. The Institutional Review Board (IRB) at the University of California, San Francisco, and Stanford University oversaw approval. Access to data followed current laws, regulations, and IRB guidelines. WSI patient samples were de-identified and did not contain any personal health information.

### Code availability

The source code for the whole slide image data preparation and convolutional neural networks, including trained model weights, is freely available under the open-source MIT license at http://github.com/keiserlab/dermato-paper.

### Author contributions

Conceptualization: E.S.K, M.J.K; Methodology: M.T., G.G., S.G., E.S.K., M.J.K.; Software: M.T., G.G., S.G., N.M.; Validation: M.T., S.G., E.S.K.; Formal analysis: M.T., G.G., S.G., E.S.K.; Investigation: M.T., G.G., S.G., N.M.; Resources: E.S.K., M.J.K.; Data Curation: M.T., G.G., S.G., N.M., E.S.K.; Writing - Original Draft: M.T., G.G., S.G., E.S.K., M.J.K.; Writing - Review & Editing: all; Visualization: M.T., G.G., S.G.; Supervision: E.S.K., M.J.K.; Project administration: M.J.K; Funding acquisition: M.J.K.

## Acknowledgments

This work was supported by CZI grant DAF2018-191905 (DOI 10.37921/550142lkcjzw) from the Chan Zuckerberg Initiative DAF, an advised fund of Silicon Valley Community Foundation (funder DOI 10.13039/100014989) (M.J.K.). We thank Dr. Kerri Rieger at Stanford University for her mentorship.

## Declaration of generative AI and AI-assisted technologies in the writing process

During the preparation of this work the authors used ChatGPT-4o (https://chat.openai.com, August 2024) to suggest edits for readability, clarity, and conciseness after we had written the manuscript draft. After using this tool/service, the authors reviewed and edited the content as needed and take full responsibility for the content of the publication.

