## Supplemental Information for "Machine-learning convergent melanocytic morphology despite noisy archival slides"

### Supplemental Figures

| Slide ID | H&E Whole Slide Image (WSI) | IHC WSI |
|---|---|---|
| WSI-01 (discarded) | 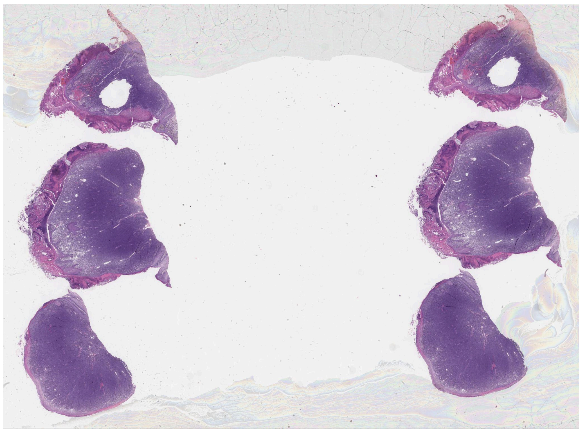 | 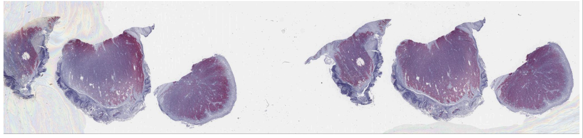 |
| WSI-02 (included) | 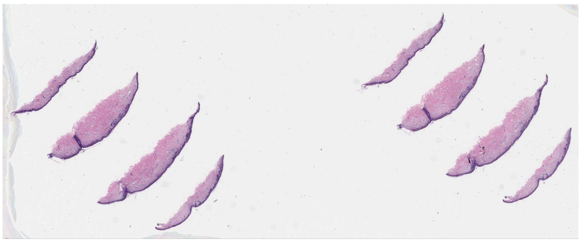 | 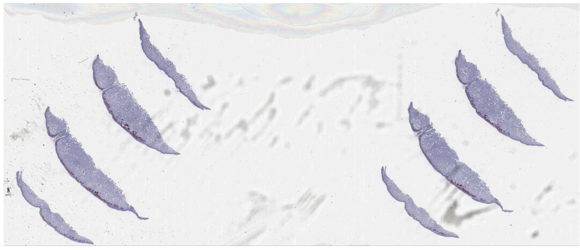 |
| WSI-03 (included) | 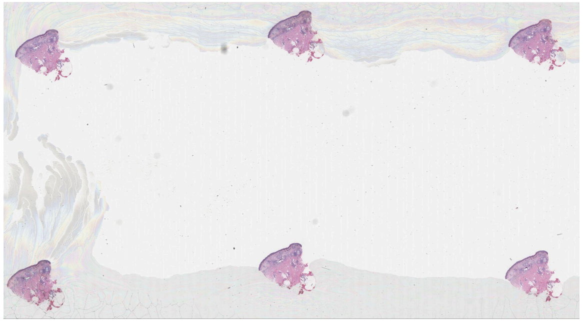 | 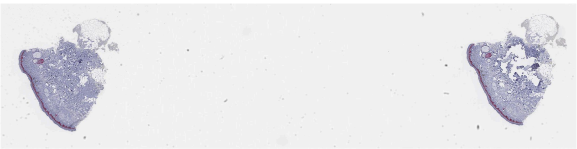 |

| | | |
|---|---|---|
| WSI-04 (discarded) | 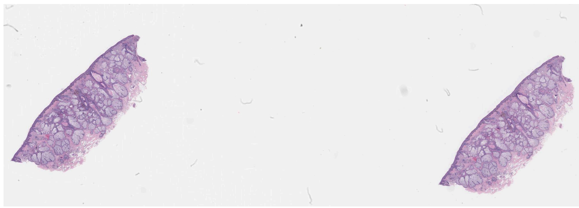 | 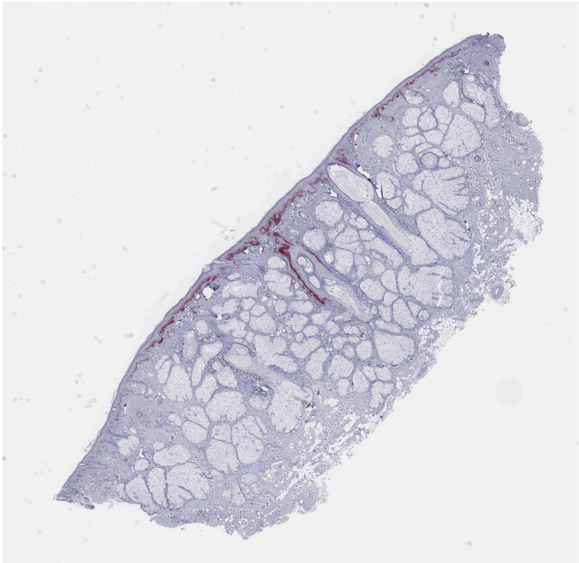 |
| WSI-05 (discarded) | 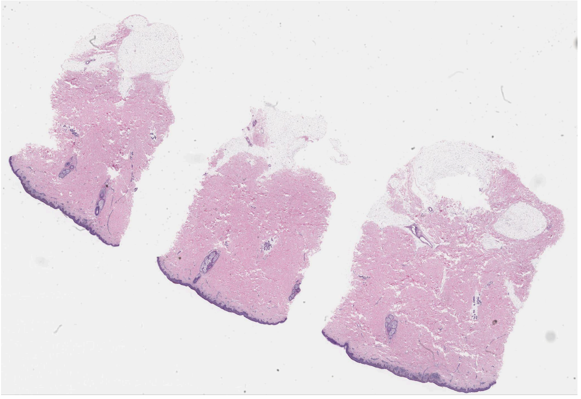 | 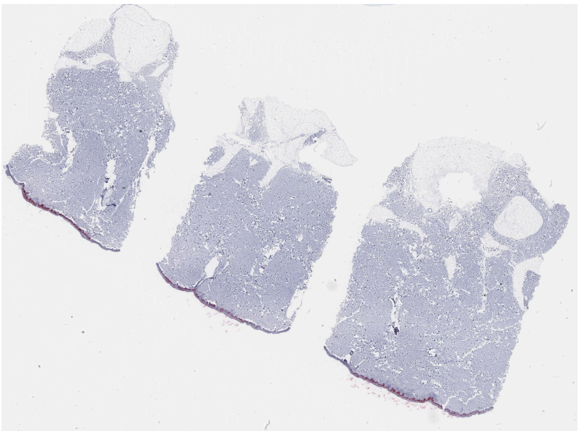 |
| WSI-06 (included) | 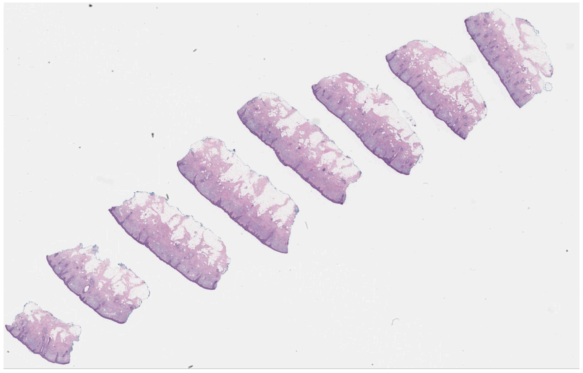 | 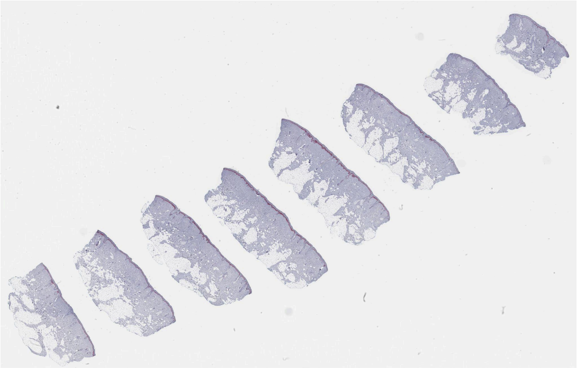 |

| | | |
|---|---|---|
| WSI-07 (included) | 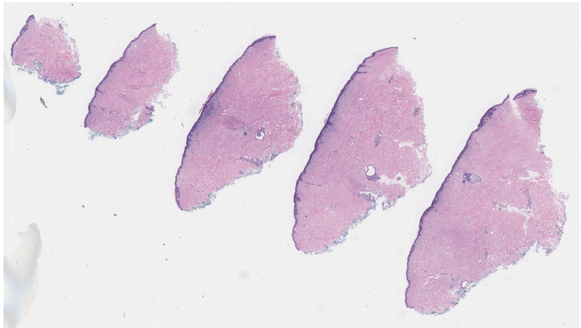 | 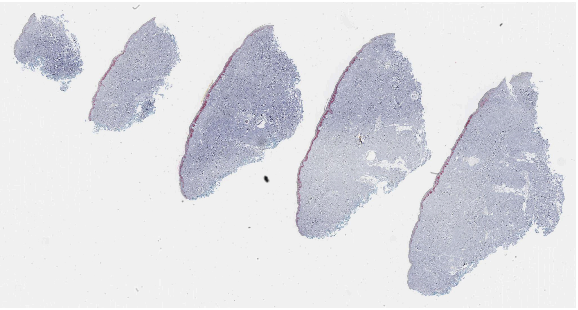 |
| WSI-08 (discarded) | 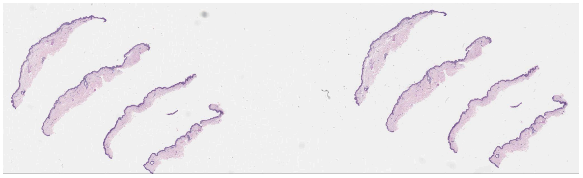 | 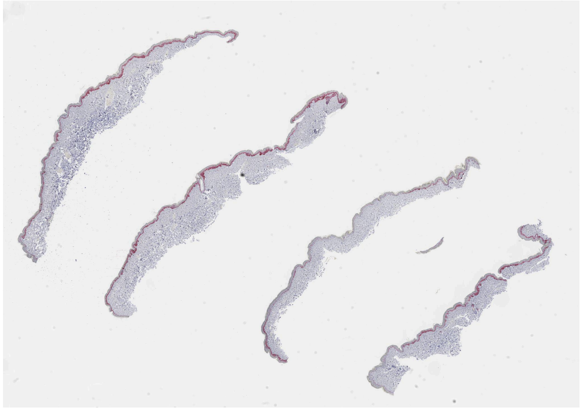 |
| WSI-09 (included) | 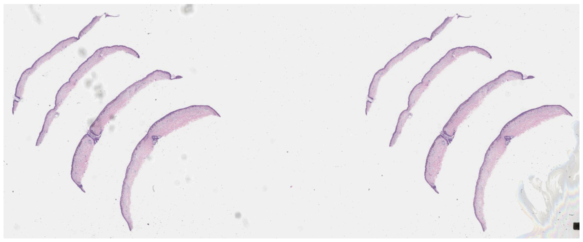 | 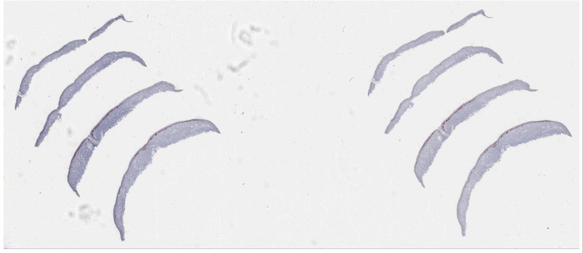 |
| WSI-10 (discarded) | 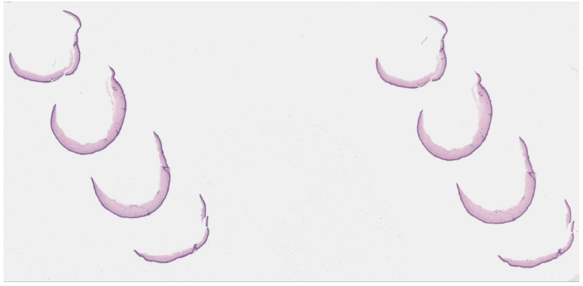 | 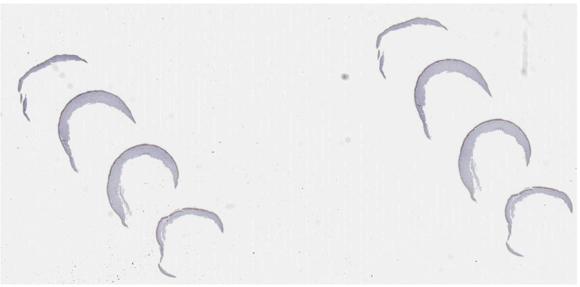 |

| | | |
|---|---|---|
| WSI-11 (discarded) | 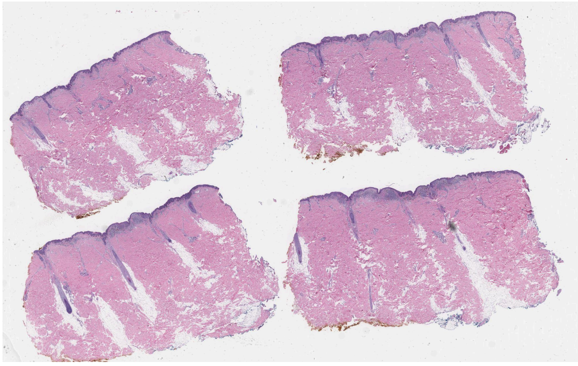 | 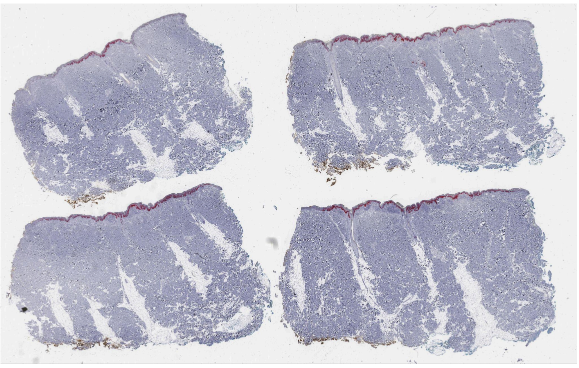 |
| WSI-12 (included) | 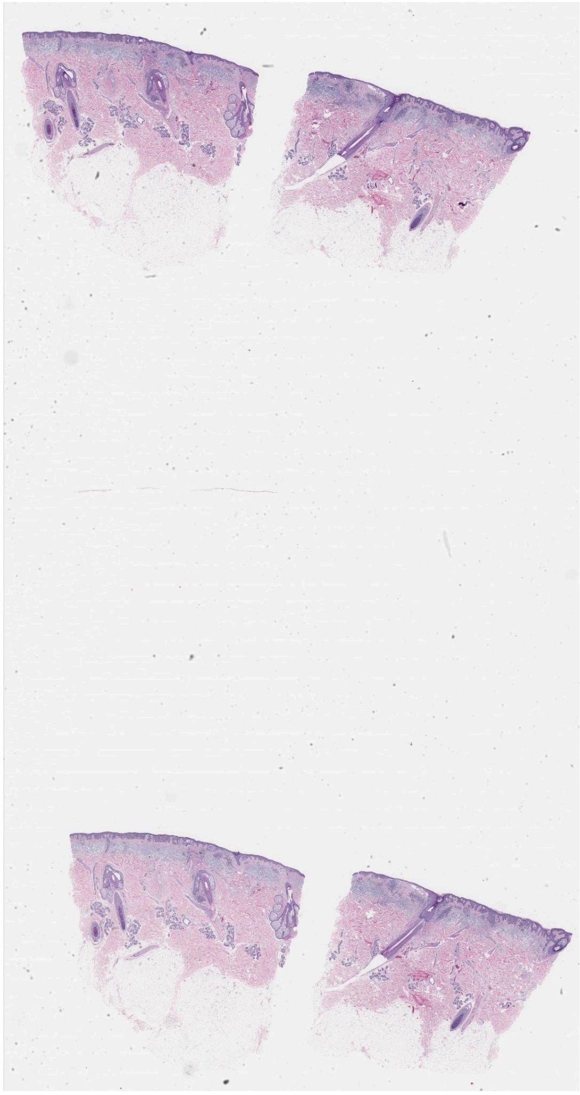 | 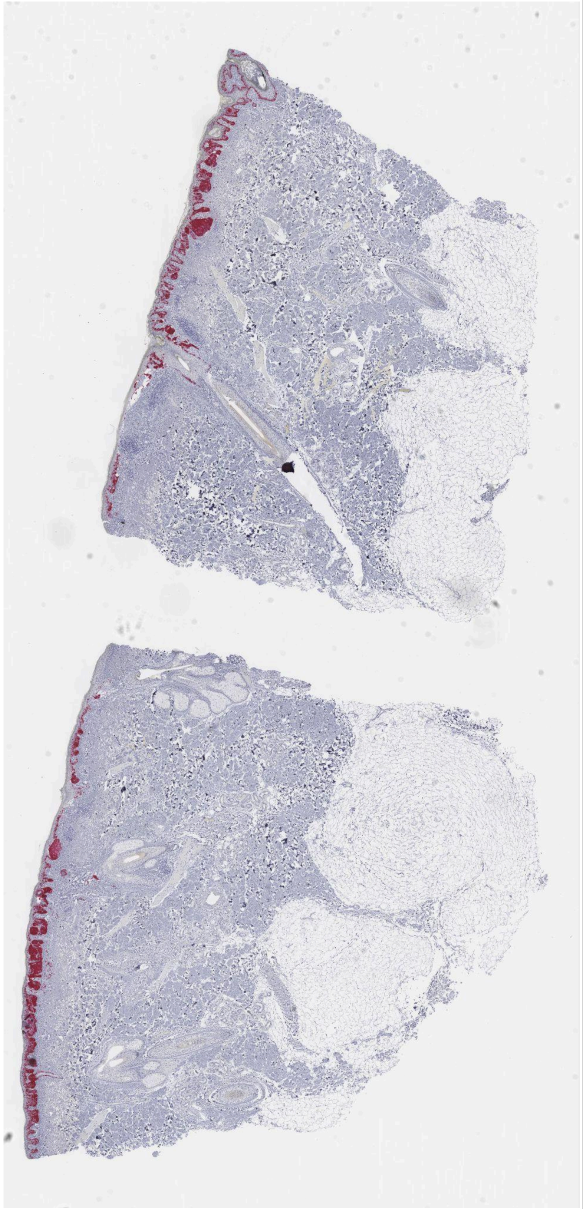 |

| | | |
|---|---|---|
| WSI-13 (included) | 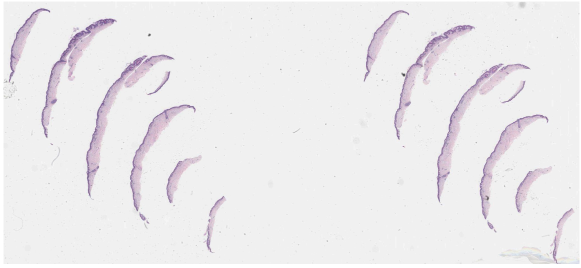 | 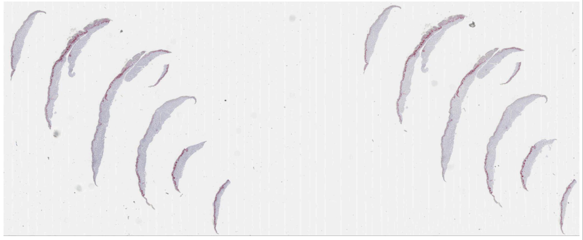 |
| WSI-14 (included) | 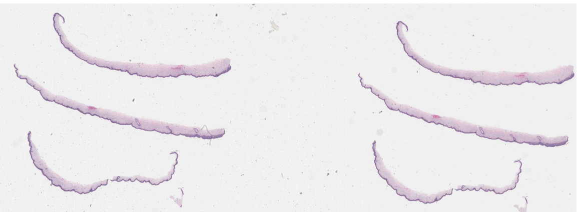 | 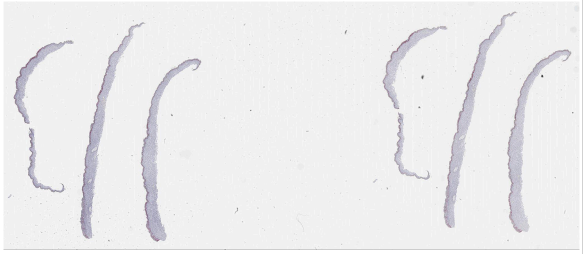 |
| WSI-15 (included) | 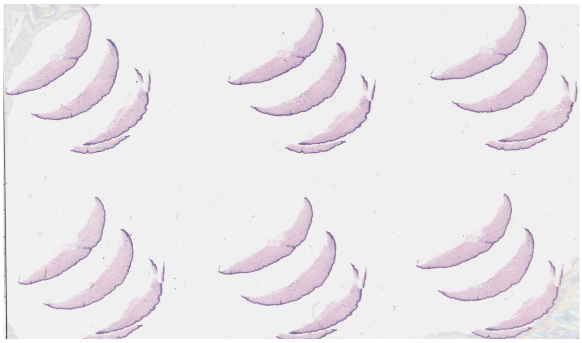 | 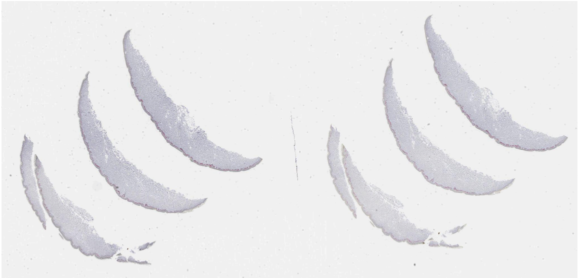 |
| WSI-16 (included) | 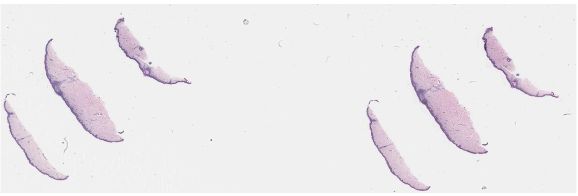 | 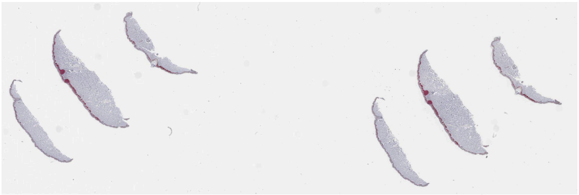 |
| WSI-17 (discarded) | 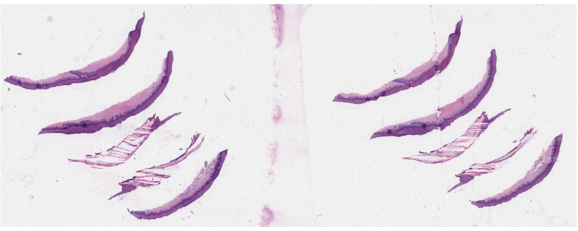 | 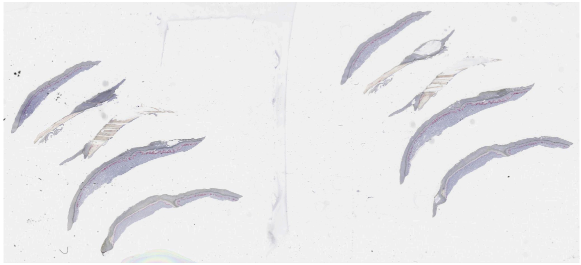 |

| | | |
|---|---|---|
| WSI-18<br>(discarded) | 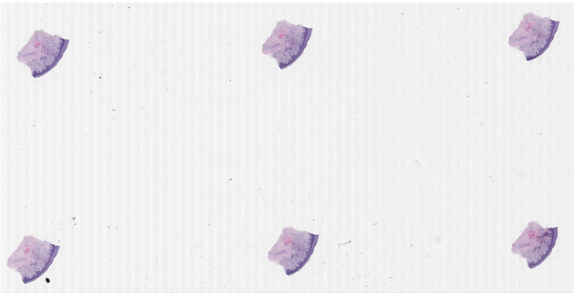 | 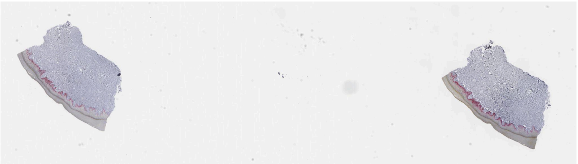 |
| WSI-19<br>(discarded) | 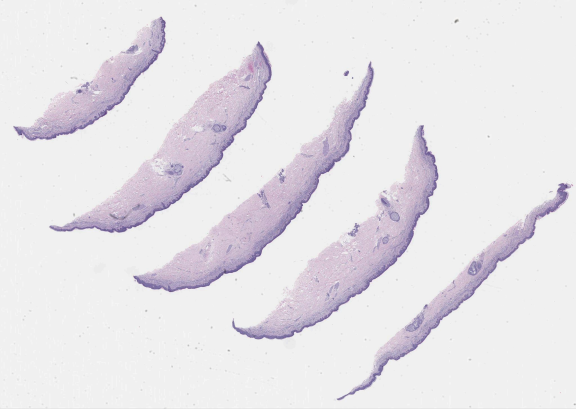 | 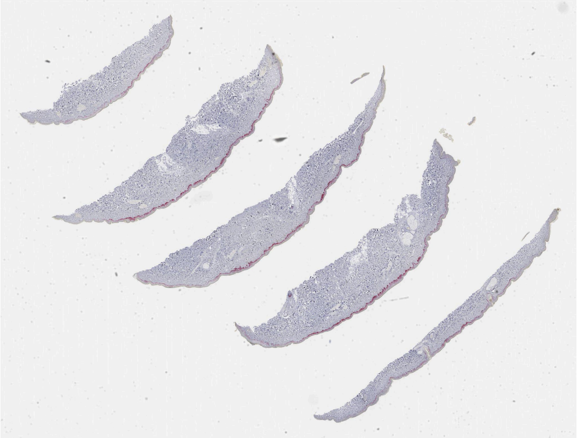 |
| WSI-20<br>(discarded) | 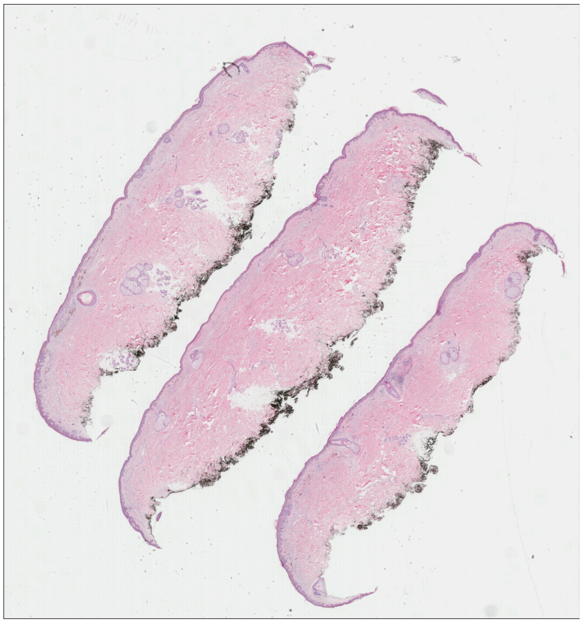 | 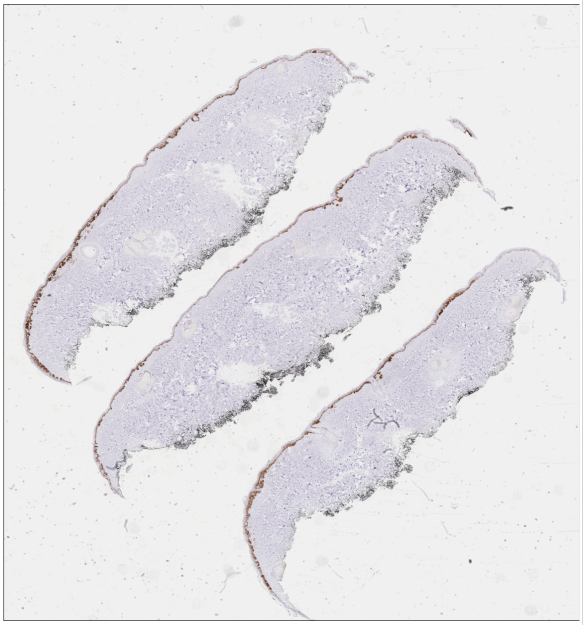 |

| | | |
|---|---|---|
| WSI-21 (discarded) | 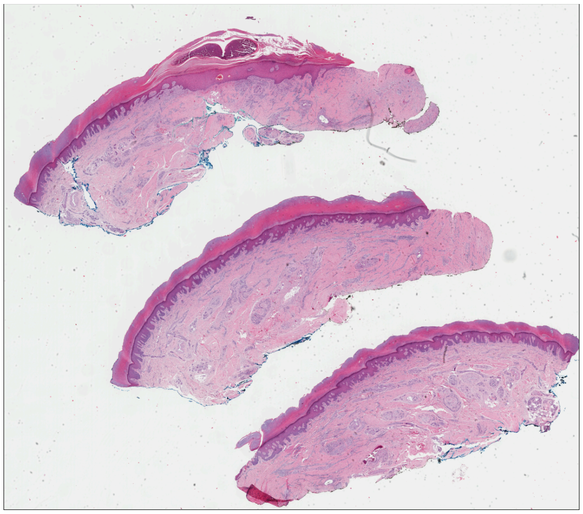 | 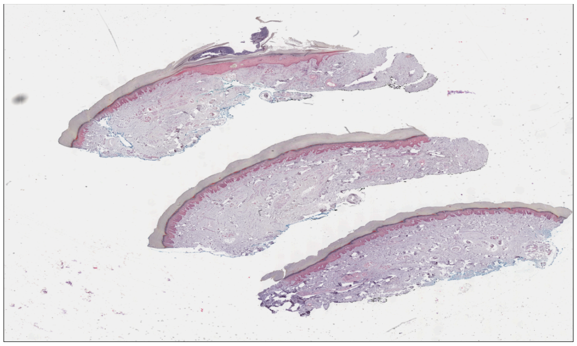 |
| WSI-22 (discarded) | 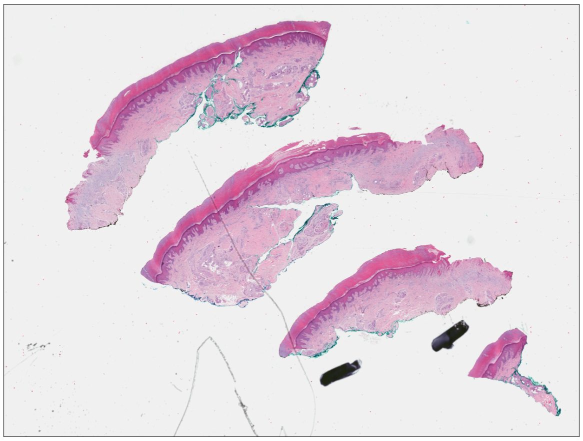 | 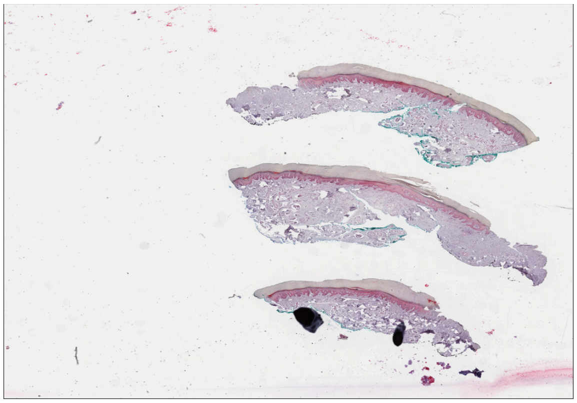 |
| WSI-23 (discarded) | 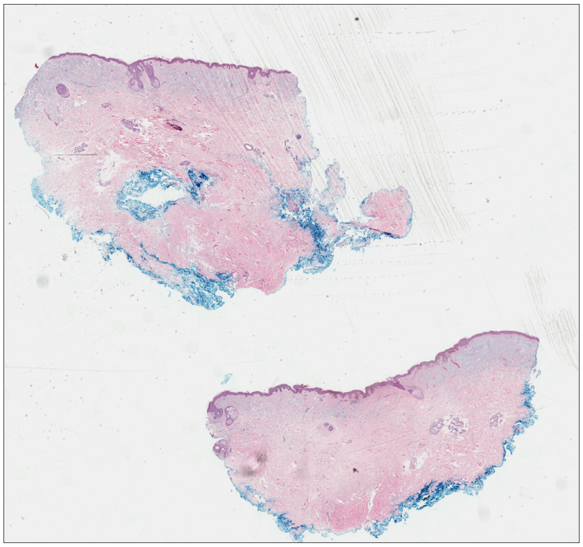 | 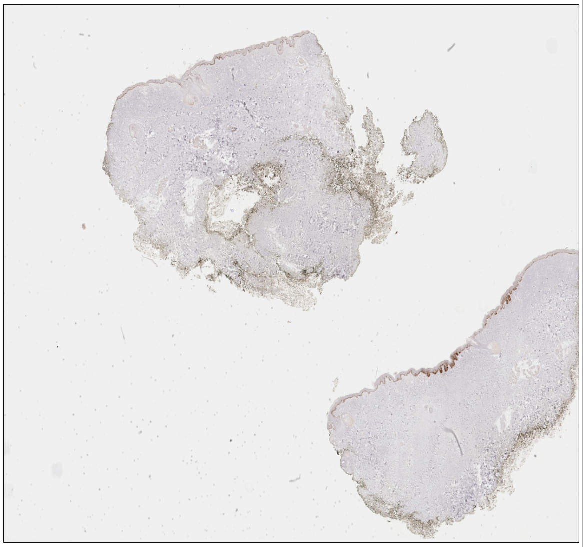 |

| | | |
|---|---|---|
| WSI-24 (discarded) | 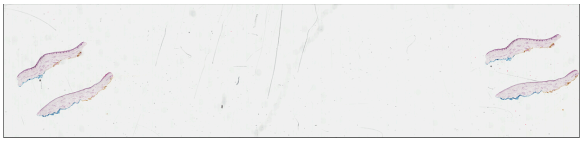 | 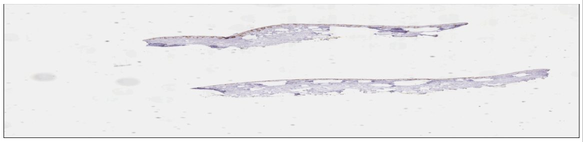 |
| WSI-25 (discarded) | 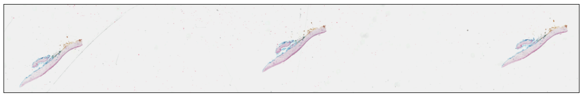 | 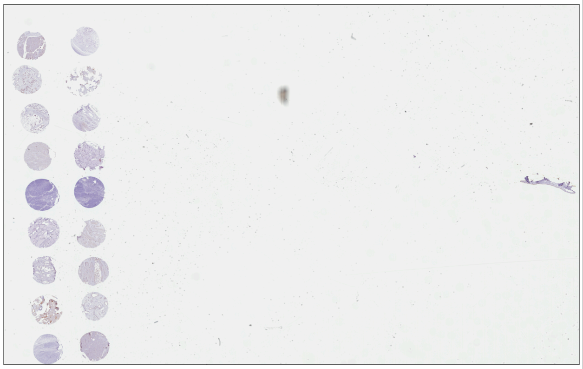 |
| WSI-26 (discarded) | 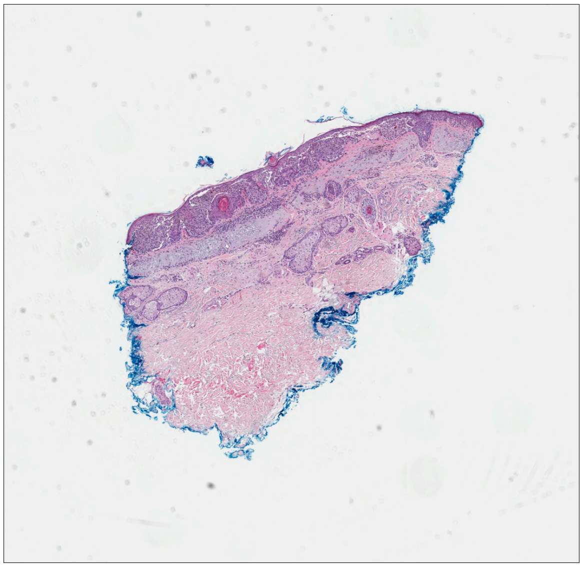 | 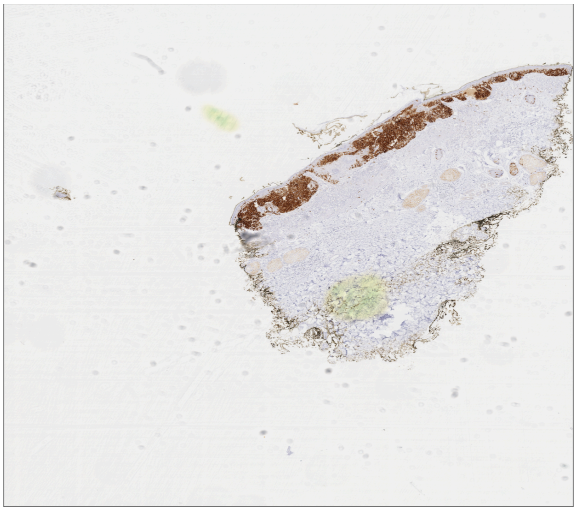 |
| WSI-27 (discarded) | 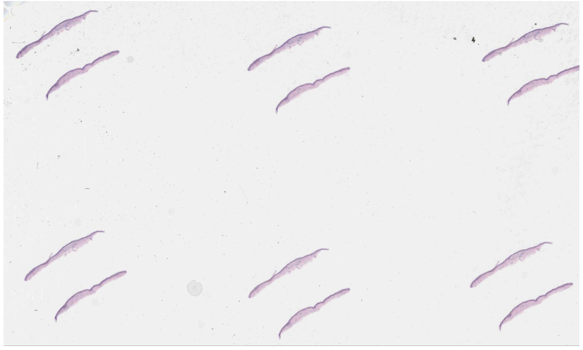 | 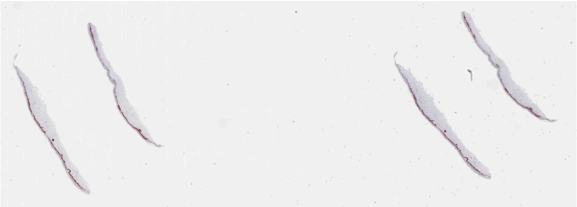 |

| | | |
|---|---|---|
| WSI-28 (included) | 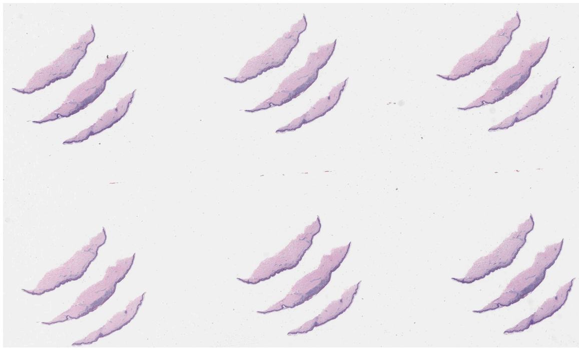 | 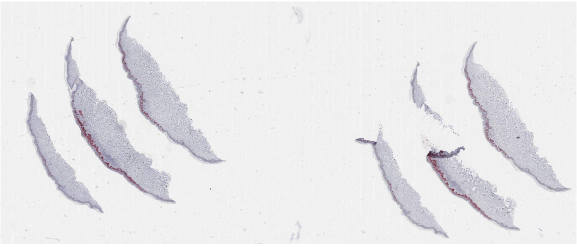 |
| WSI-29 (discarded) | 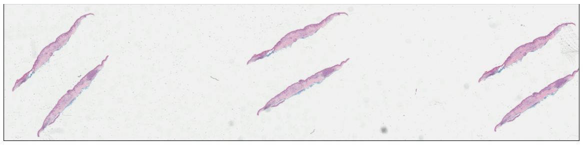 | 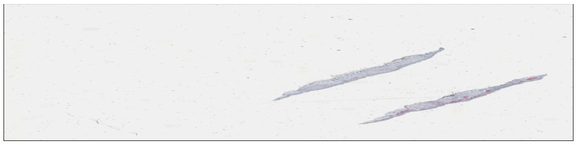 |
| WSI-30 (included) | 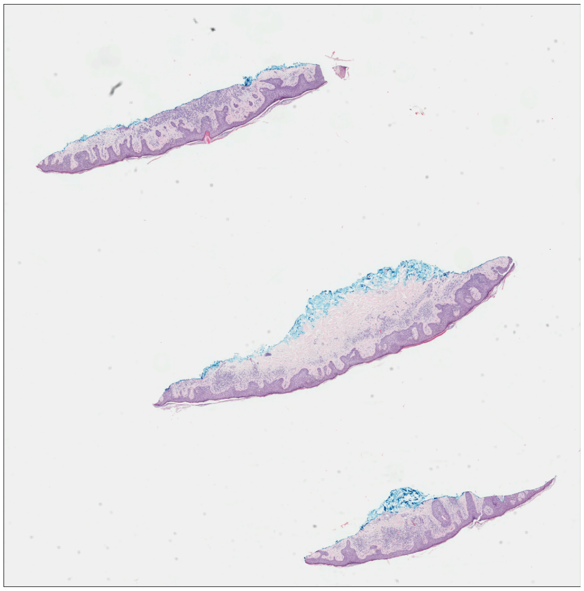 | 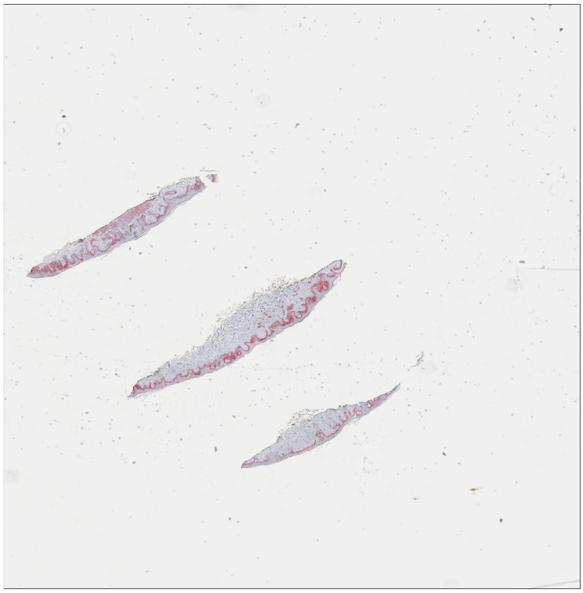 |

| | | |
|---|---|---|
| WSI-31 (discarded) | 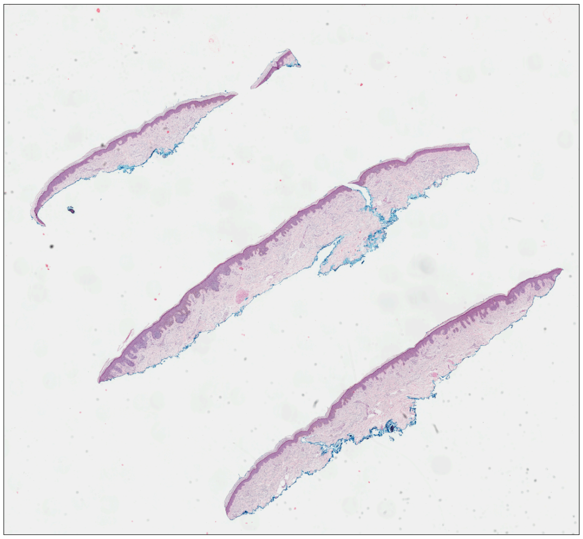 | 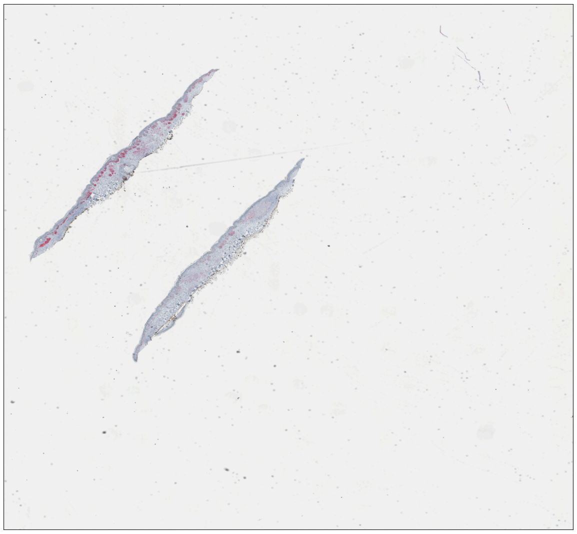 |
| WSI-32 (included) | 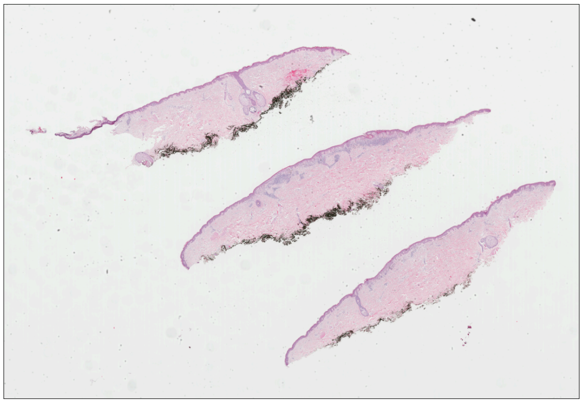 | 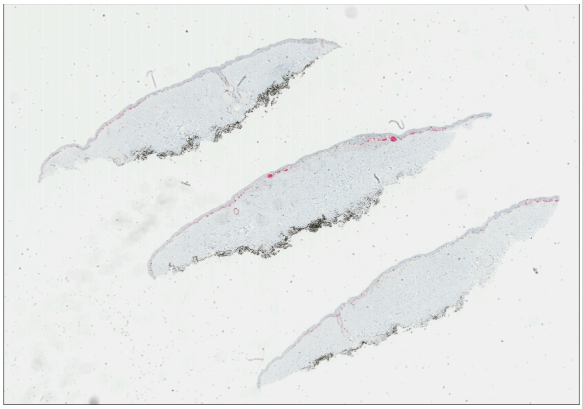 |
| WSI-33 (discarded) | 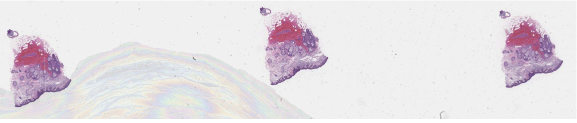 | 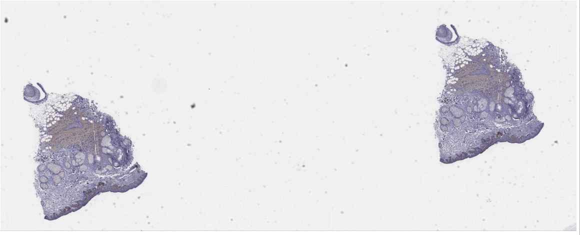 |
| WSI-34 (discarded) | 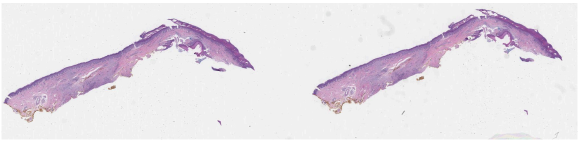 | 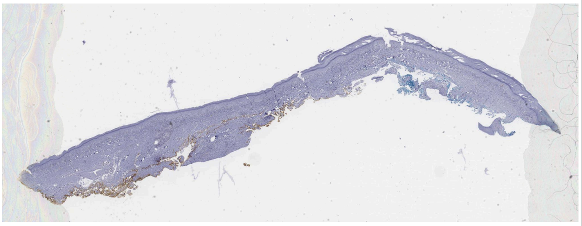 |

| | | |
|---|---|---|
| WSI-35 (discarded) | 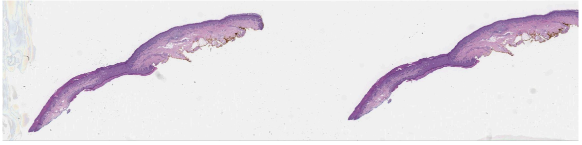 | 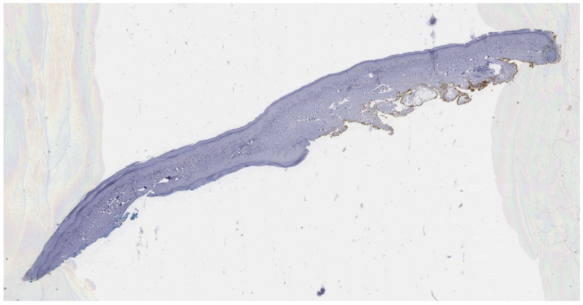 |
| WSI-36 (discarded) | 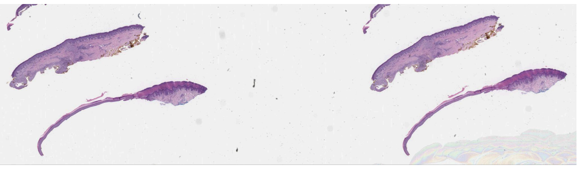 | 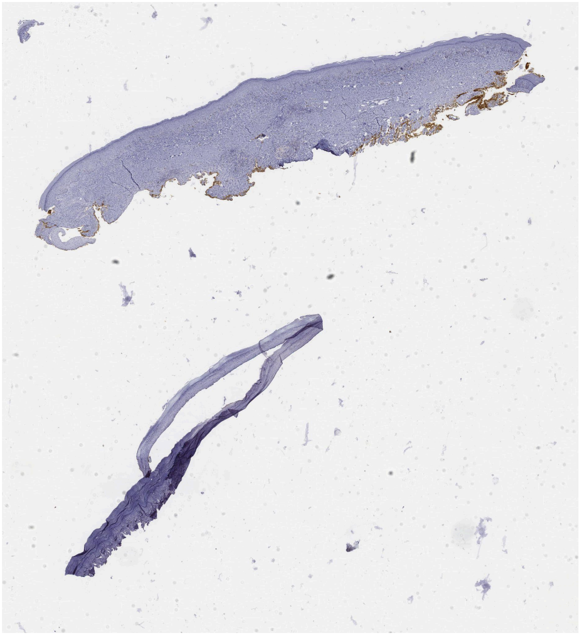 |
| WSI-37 (discarded) | 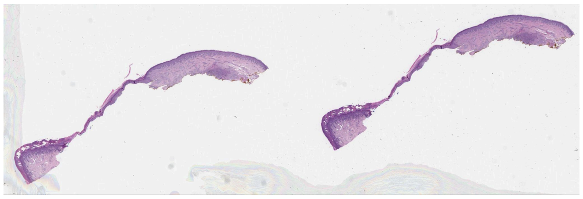 | 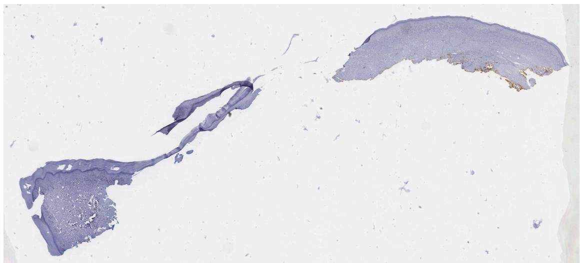 |
| WSI-38 (discarded) | 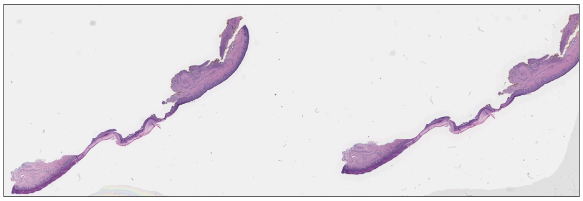 | 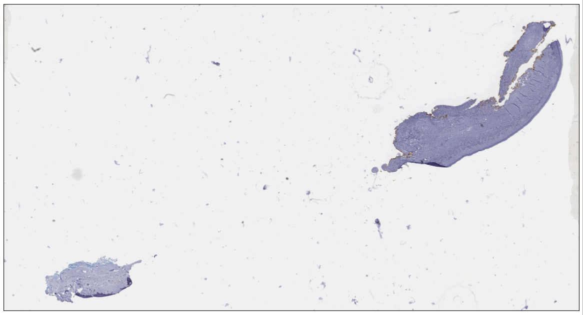 |

| | | |
|---|---|---|
| WSI-39 (discarded) | 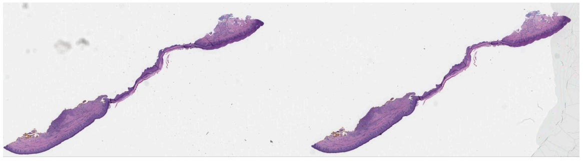 | 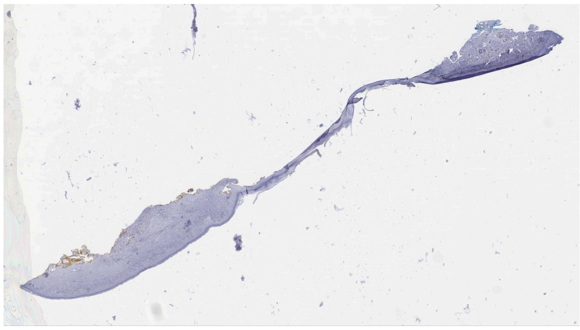 |
| WSI-40 (discarded) | 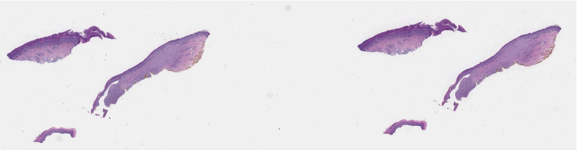 | 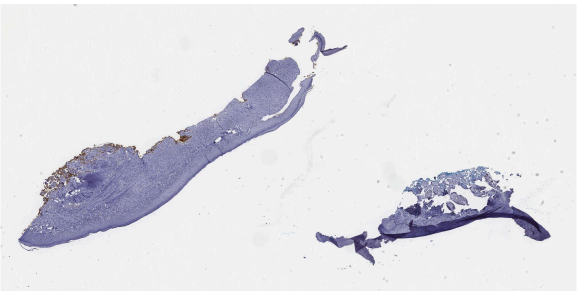 |
| WSI-41 (discarded) | 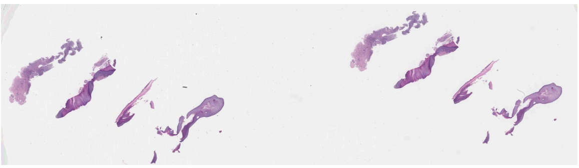 | 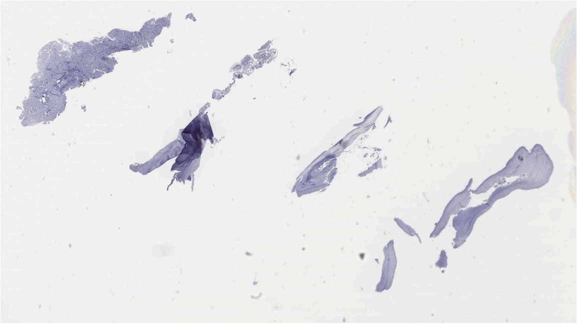 |
| WSI-42 (included) | 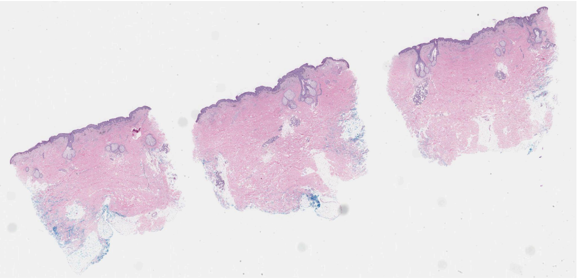 | 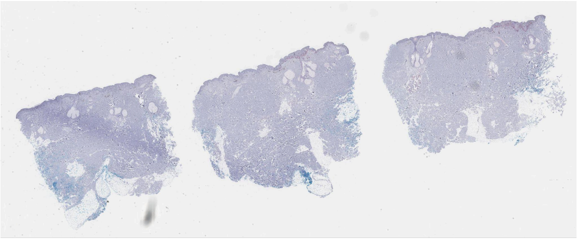 |

| | | |
|---|---|---|
| WSI-43 (discarded) | 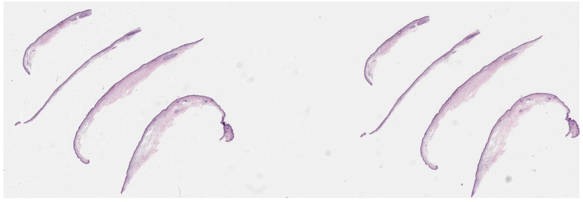 | 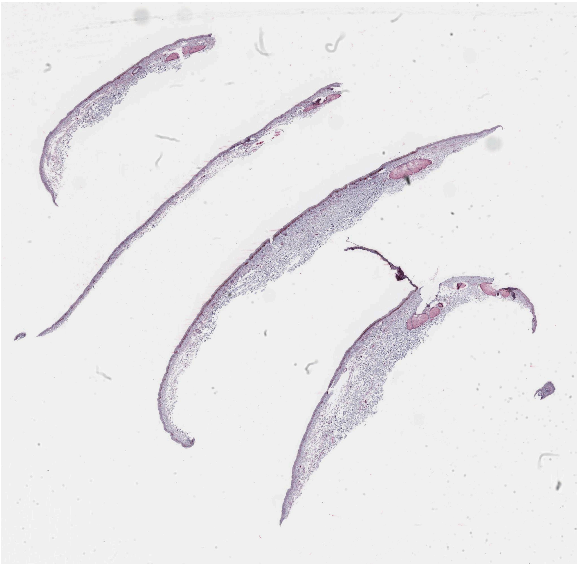 |
| WSI-44 (included) | 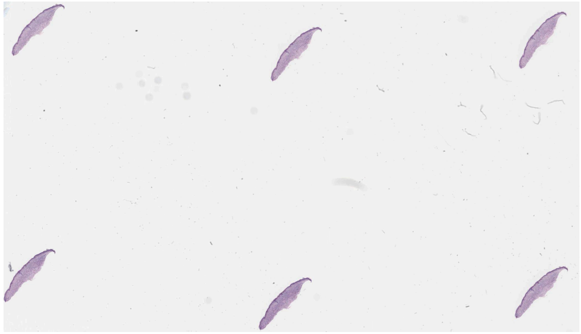 | 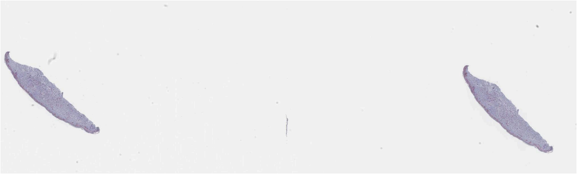 |
| WSI-45 (discarded) | 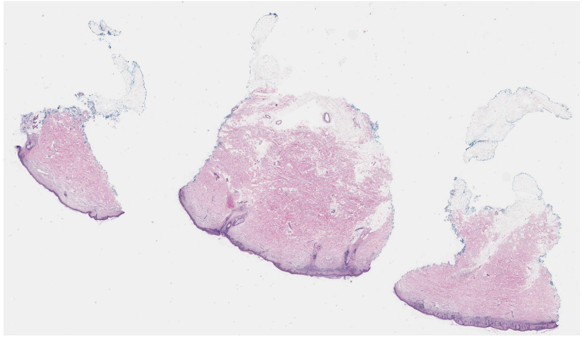 | 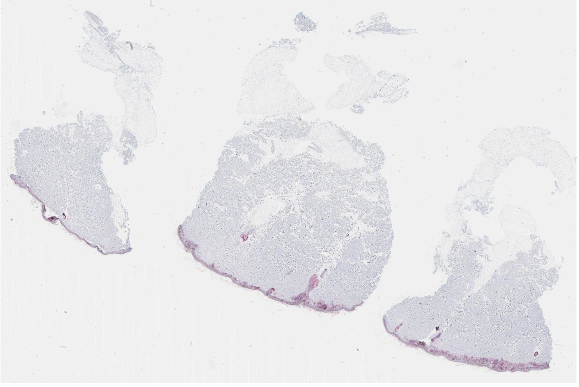 |

| | | |
|---|---|---|
| WSI-46 (discarded) | 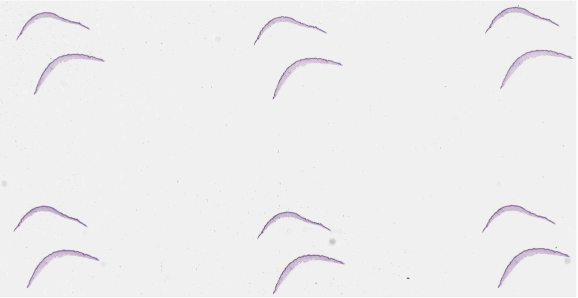 | 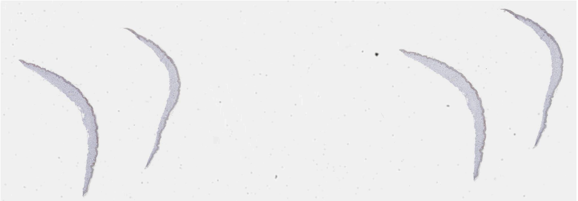 |
| WSI-47 (included) | 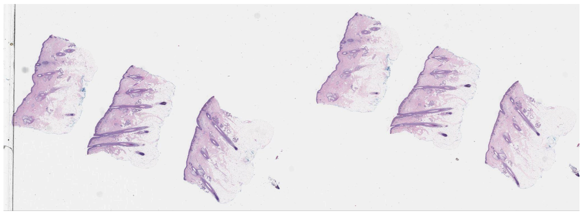 | 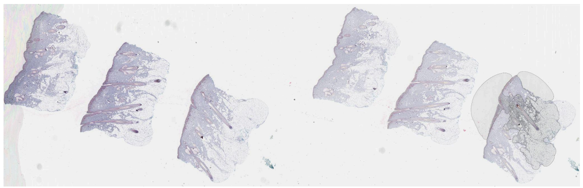 |
| WSI-48 (discarded) | 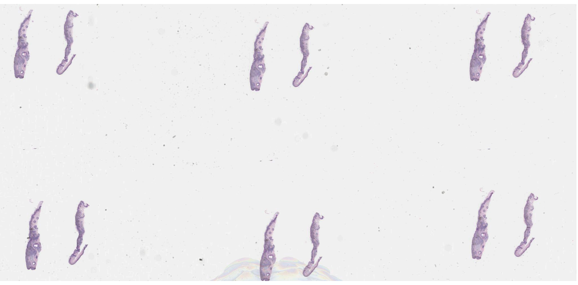 | 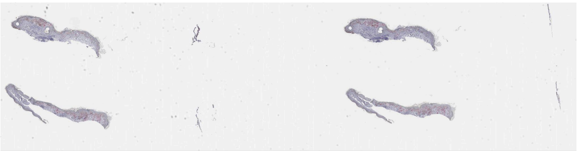 |
| WSI-49 (discarded) | 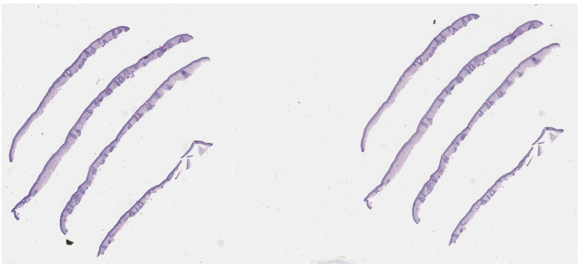 | 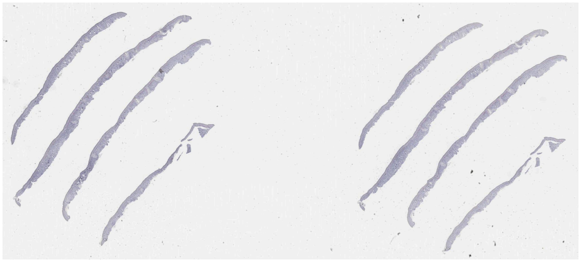 |
| WSI-50 (discarded) | 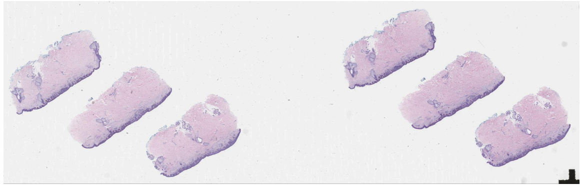 | 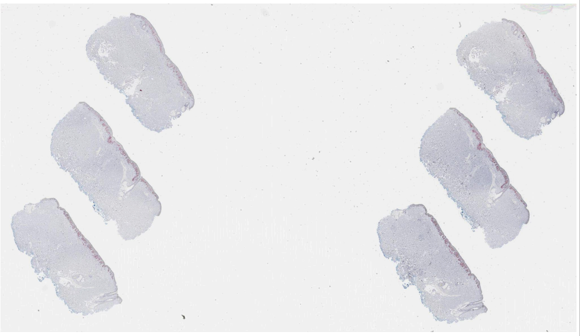 |

| | | |
|---|---|---|
| WSI-51 (discarded) | 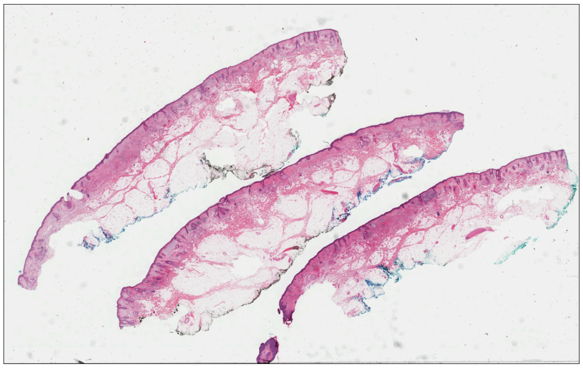 | 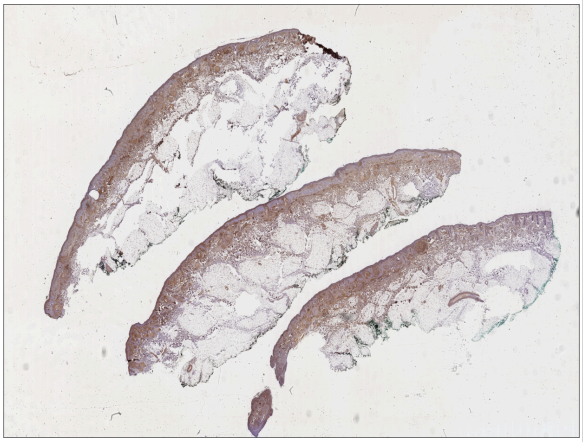 |
| WSI-52 (included) | 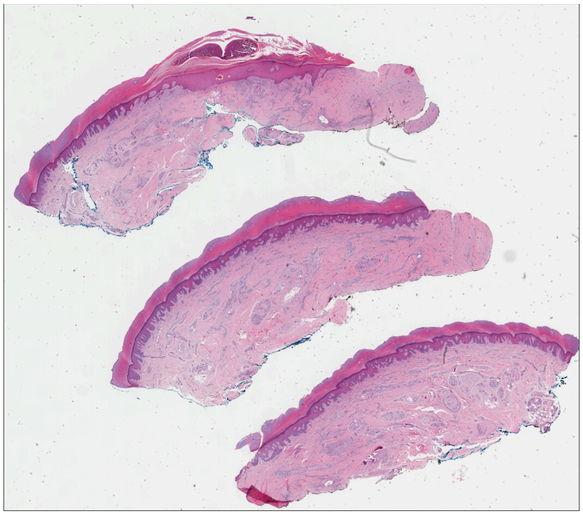 | 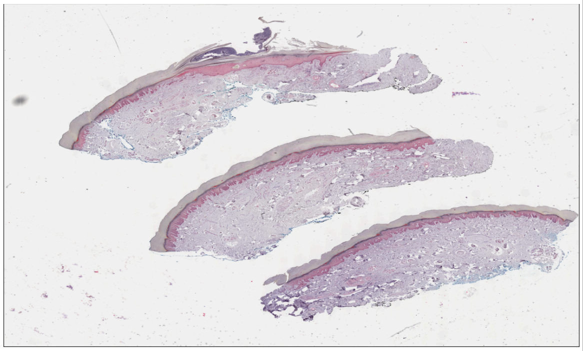 |
| WSI-53 (discarded) | 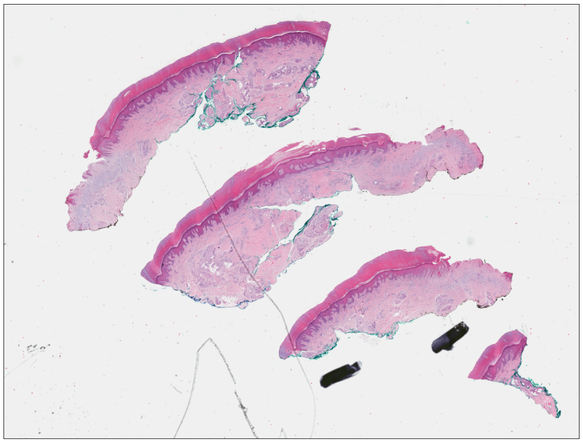 | 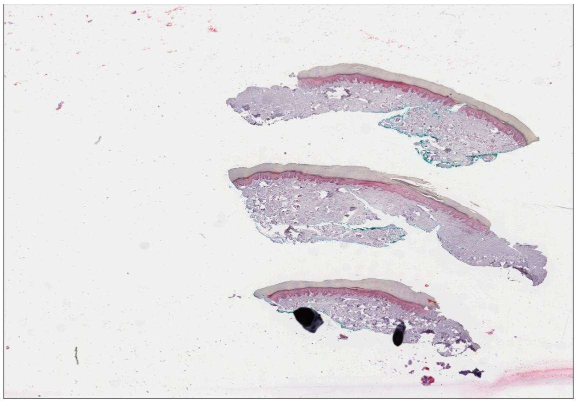 |

| | | |
|---|---|---|
| WSI-54 (discarded) | 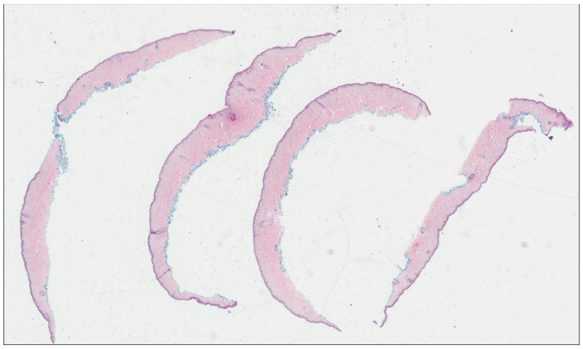 | 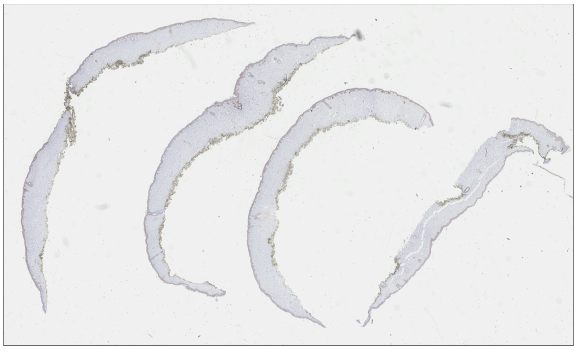 |
| WSI-55 (included) | 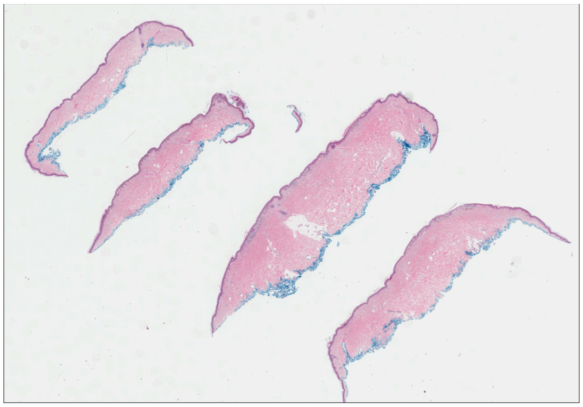 | 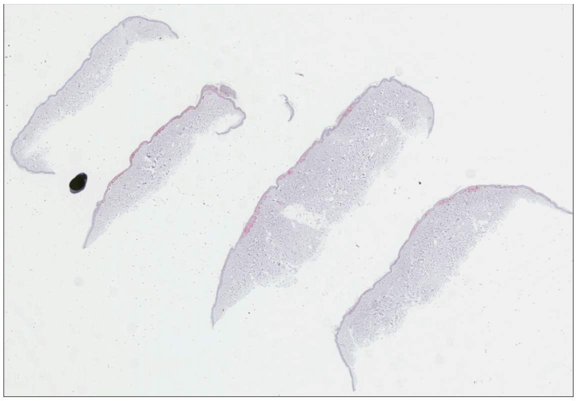 |
| WSI-56 (discarded) | 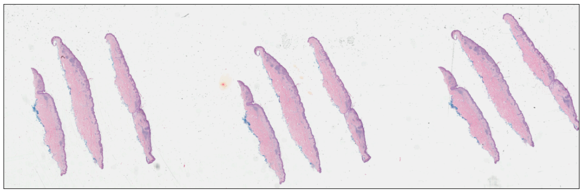 | 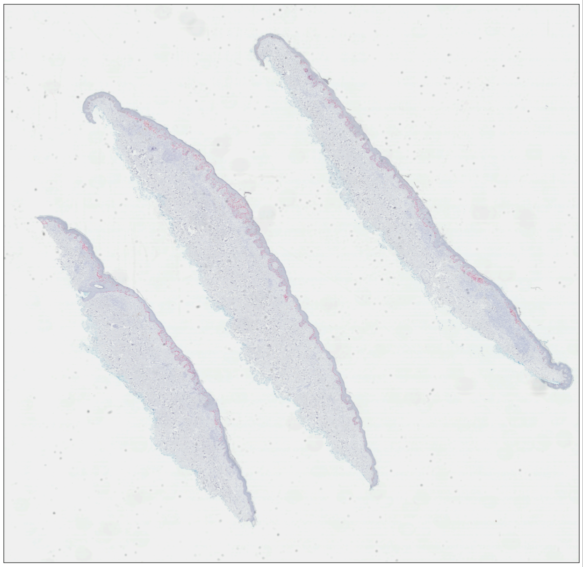 |

| | | |
|---|---|---|
| WSI-57 (included) | 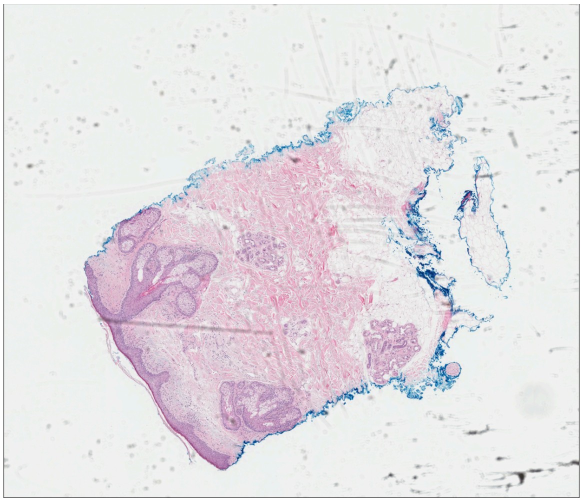 | 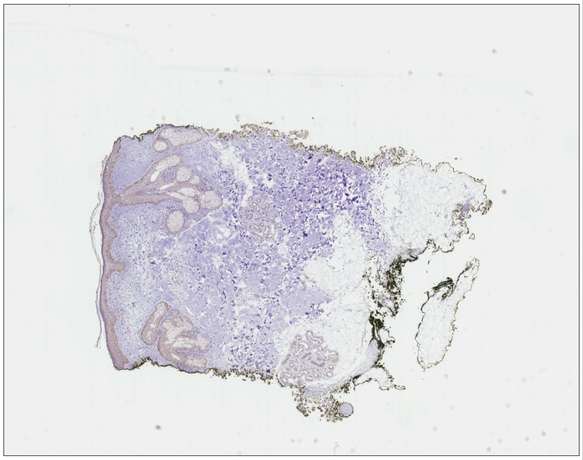 |
| WSI-58 (included) | 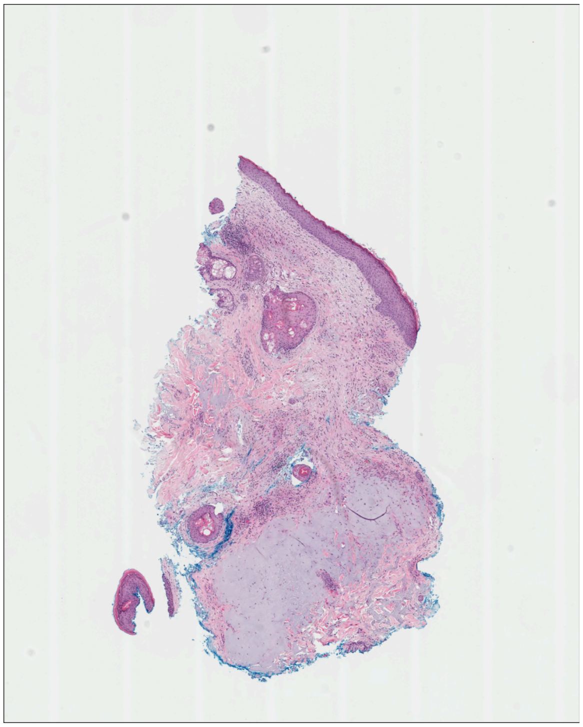 | 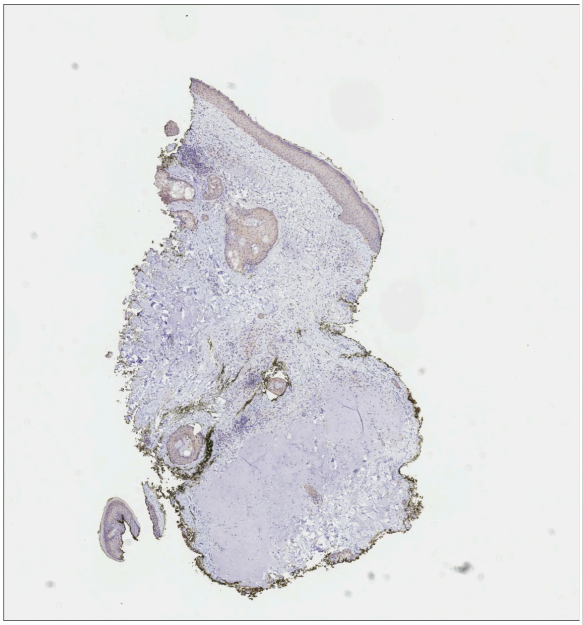 |
| WSI-59 (included) | 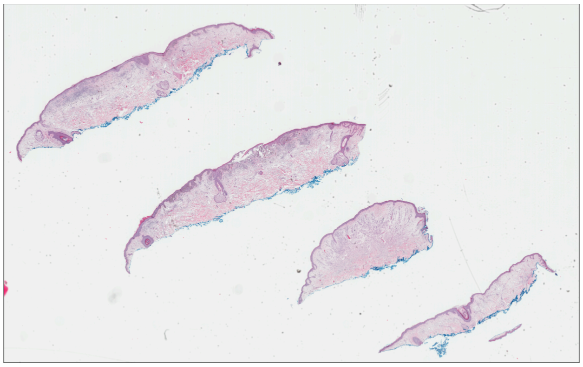 | 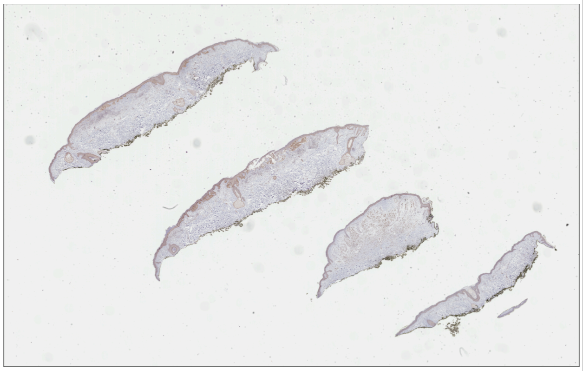 |

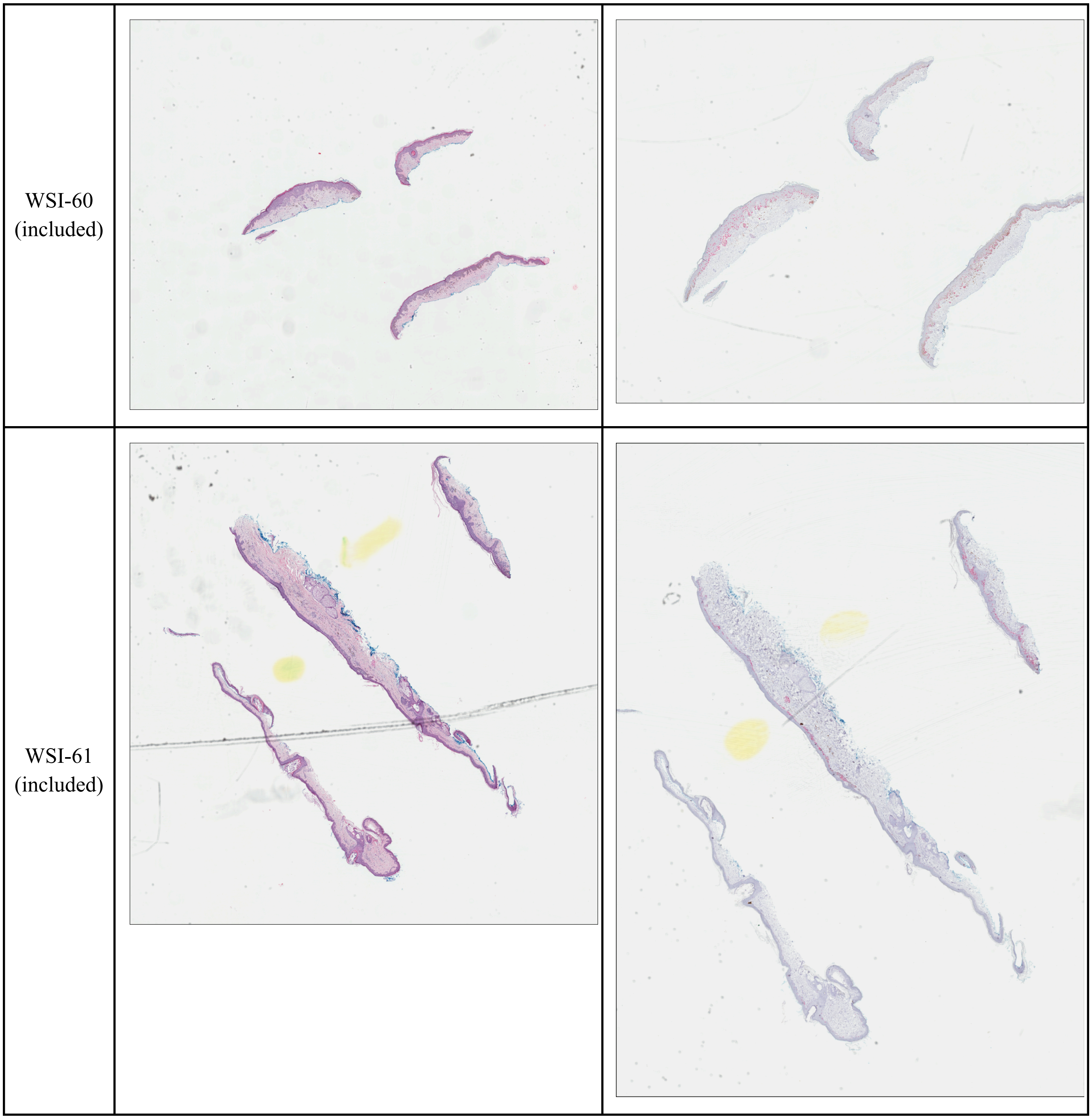


**Supplemental Figure 1.** Illustrative low-resolution summary views of archival H&E - IHC whole slide image pairs. Details are available in Supplemental Tables 1-3.

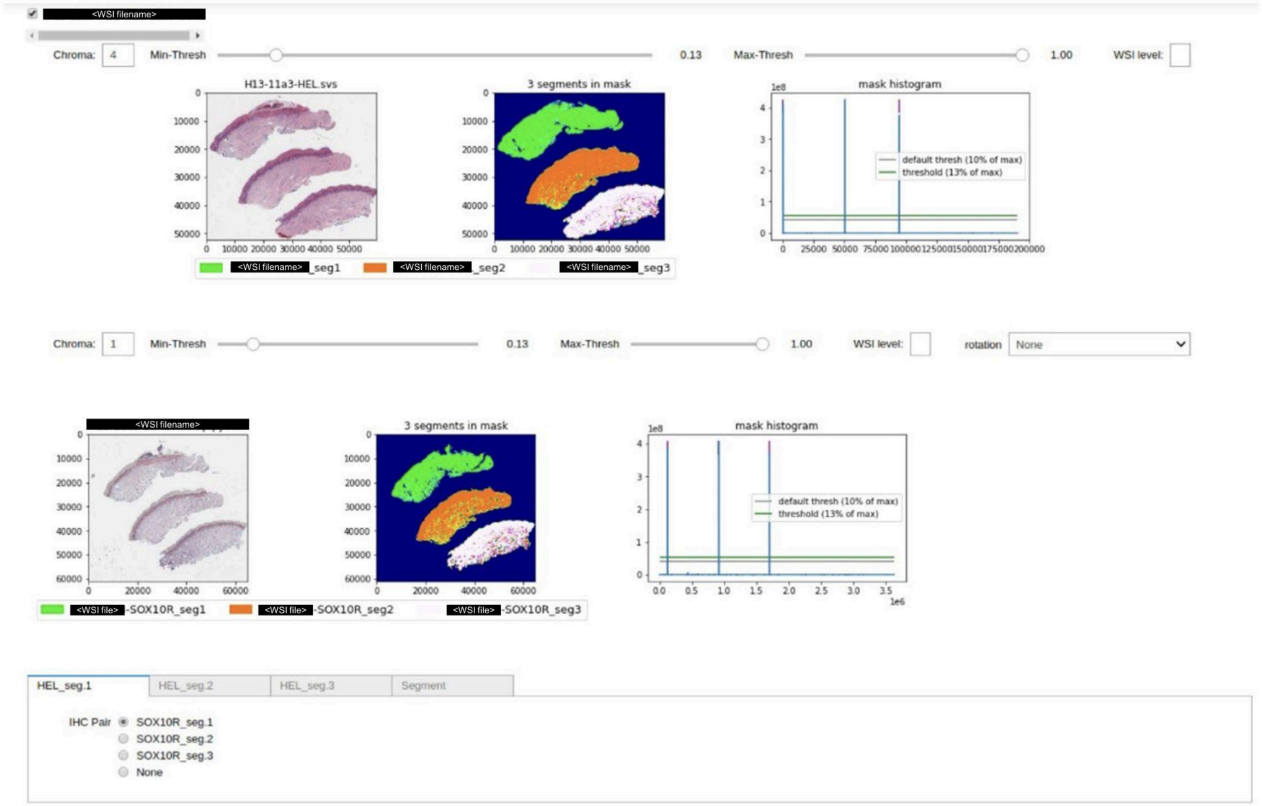


**Supplemental Figure 2**. Automated tissue extraction toolkit. An example of a sample pair extracted using the automated toolkit. The toolkit visualizes both the H&E (top, left) and IHC (middle, left) whole slide images (WSIs) and extracts tissue regions according to threshold values set for each of the ipywidgets provided. Matching occurs automatically based on tissue locations in the WSI; the user can manually adjust it if necessary (bottom).

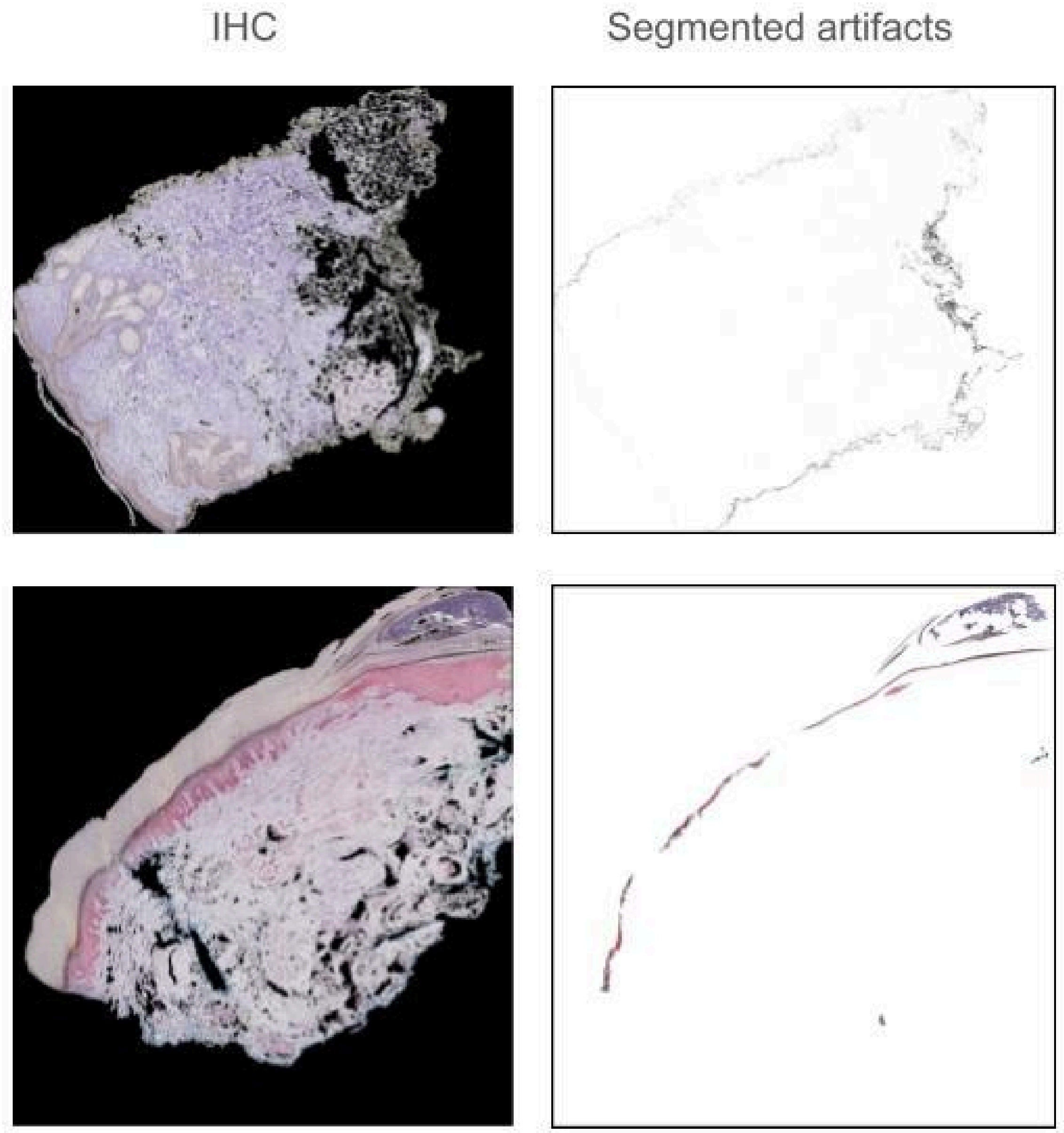


**Supplemental Figure 3.** Examples of artifacts that we computationally removed from IHC WSIs. The top row shows ink, and the bottom shows blood.

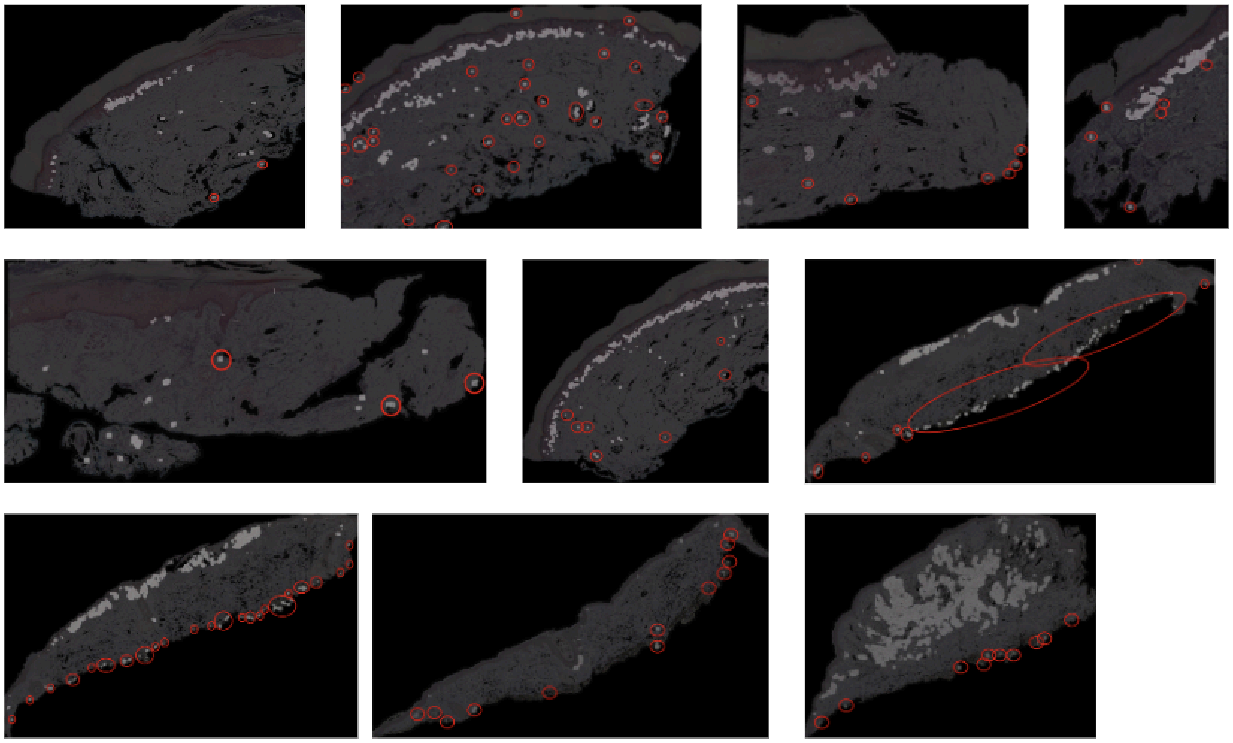

**Supplemental Figure 4.** Manual label correction by removal of false-positive areas. The labeling pipeline sometimes misidentified non-melanocyte regions as melanocytes due to tissue artifacts. To ensure label accuracy, we manually corrected these errors (red circles).

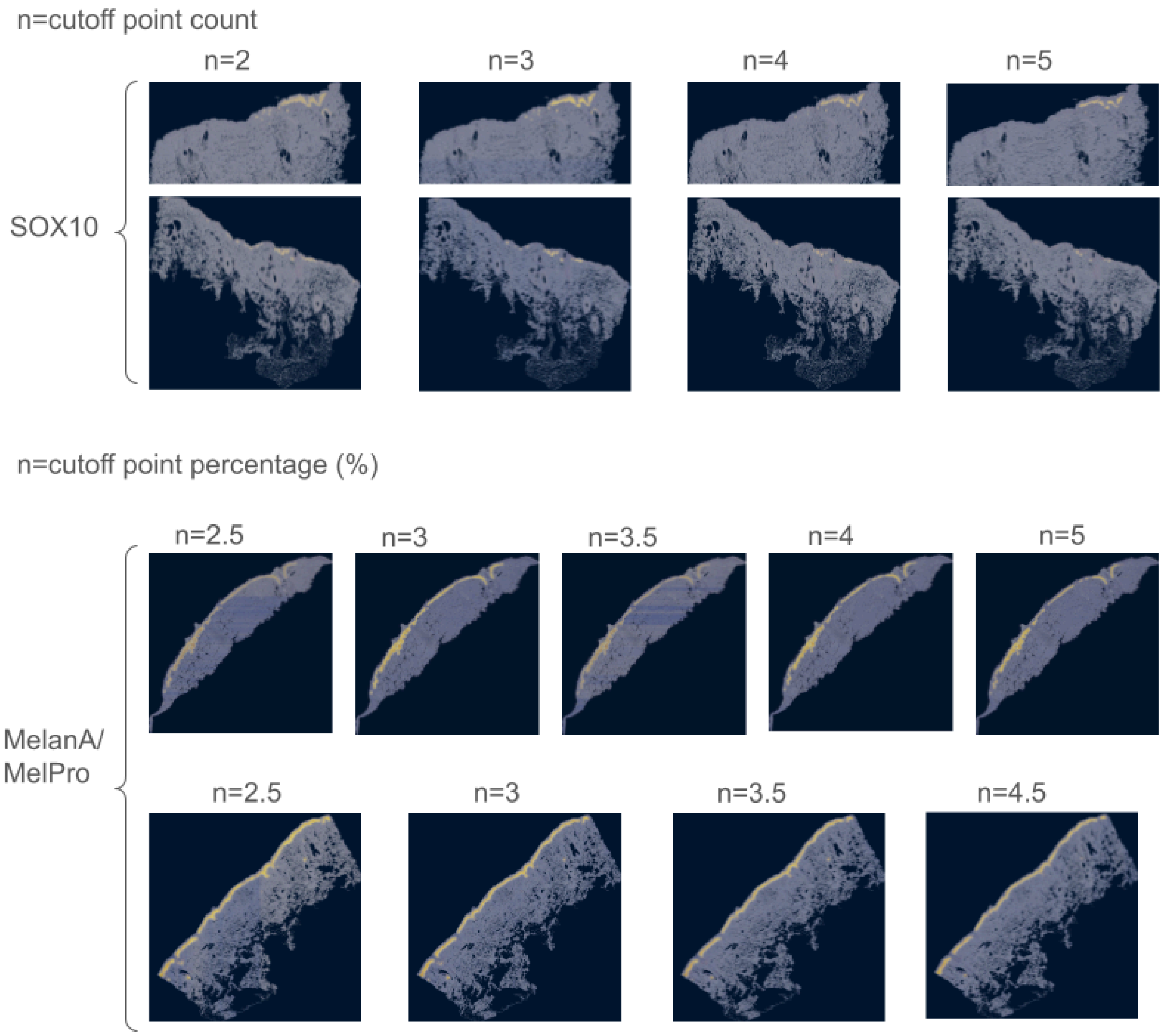


**Supplemental Figure 5.** “Label map” examples using progressively increasing (more stringent) minimum-signal thresholds, *n*, for two SOX10 (top row) and two MelanA/MelPro (bottom row) WSIs. Label maps derive directly from positive IHC stain but are coarser-grained than the stain itself due to the 256x256-pixel tiles.

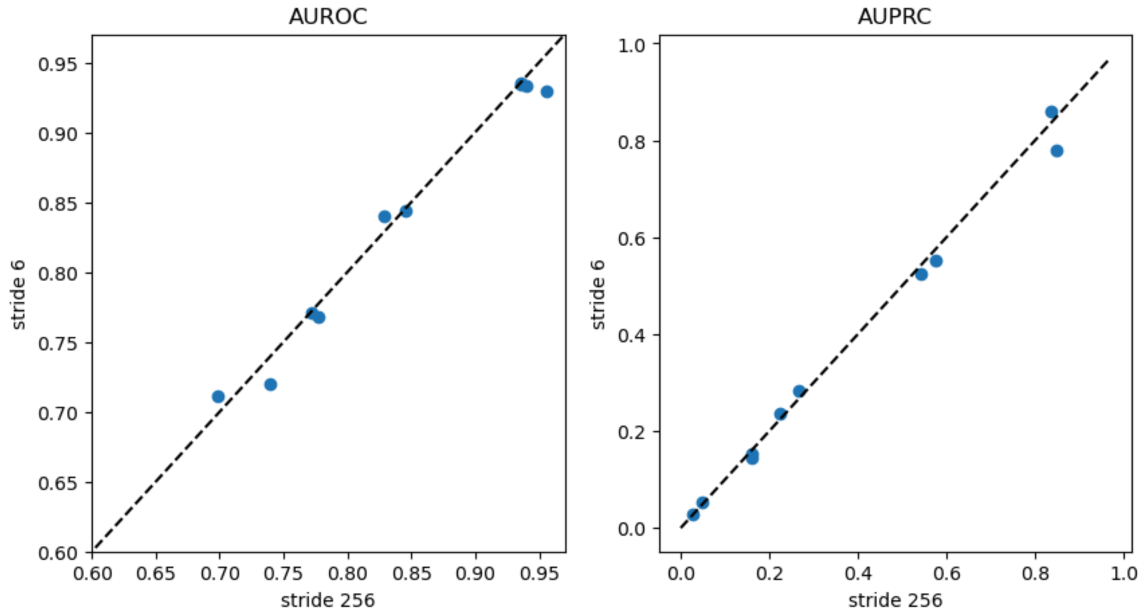


**Supplemental Figure 6.** Comparison of performance calculated by AUROC score using stride 6-pixel (y-axis) versus 256-pixel (x-axis) strides on two test sets. Each blue dot represents a tissue section. Stride 256-pixel calculations are much faster to compute due to the $n^2$ scaling of the heatmap calculation (where $n$ is proportional to 1/stride).

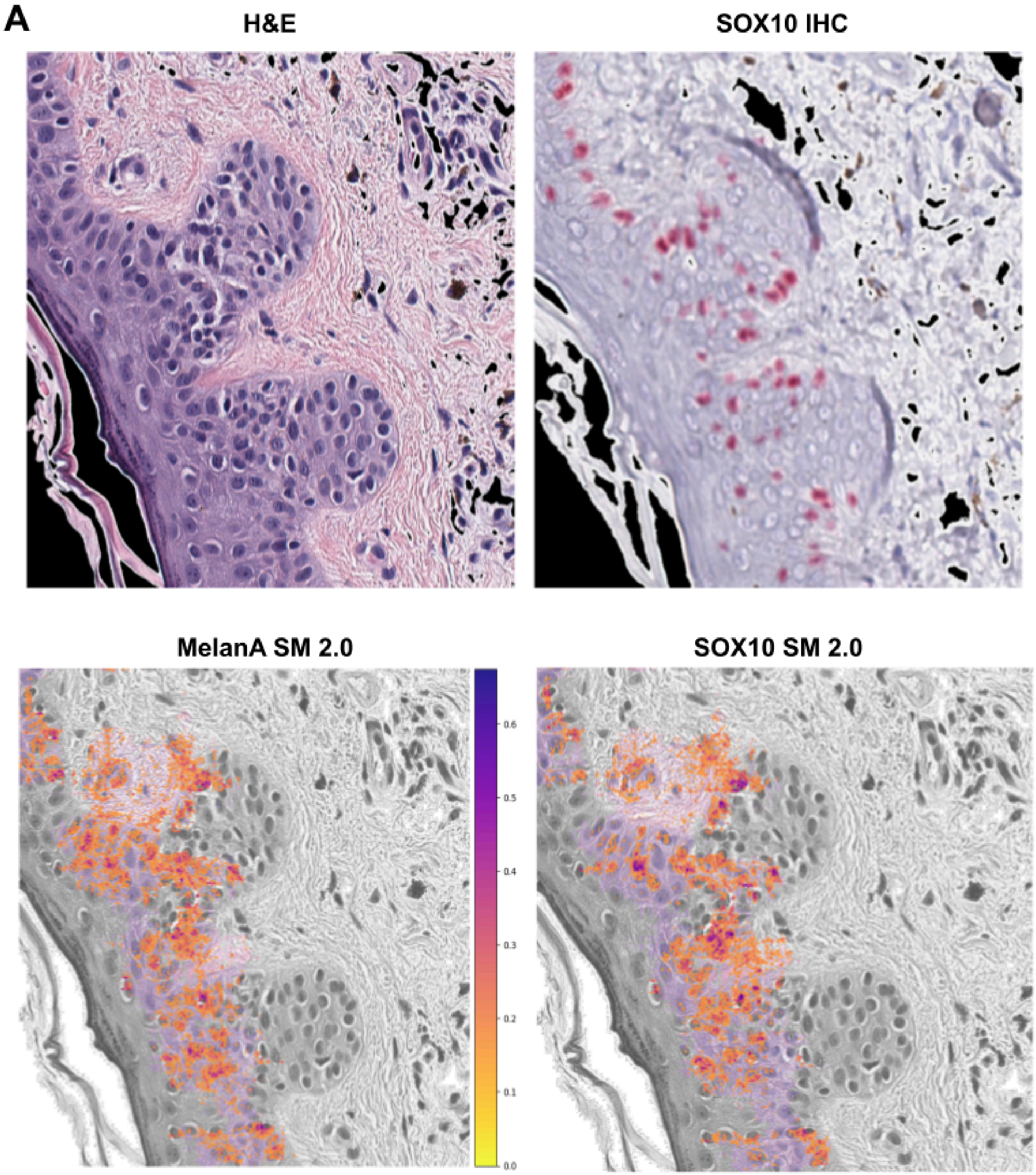
A
H&E
SOX10 IHC
MelanA SM 2.0
SOX10 SM 2.0
0.6
0.5
0.4
0.3
0.2
0.1
0.0

B

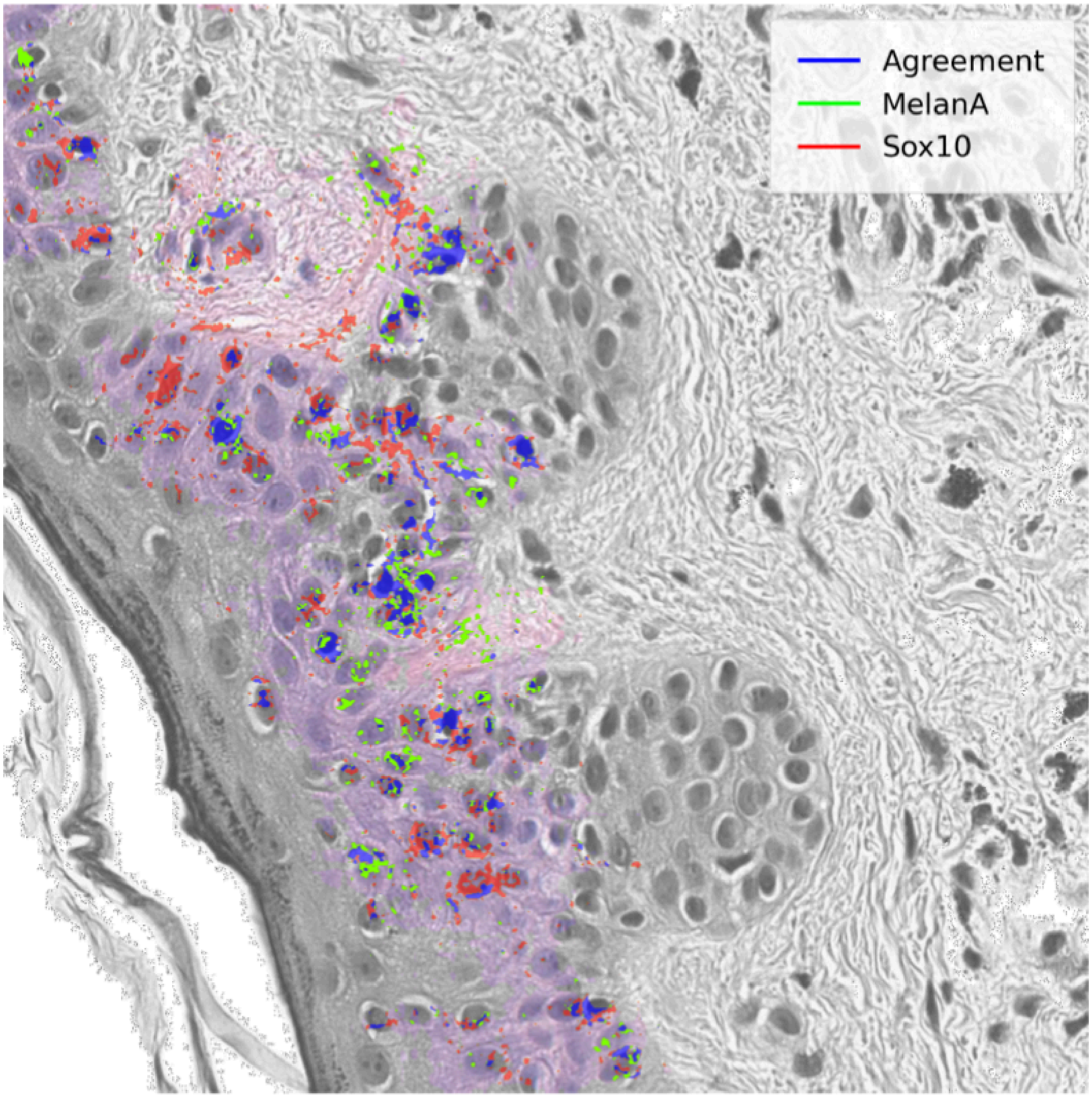


**Supplemental Figure 7. Recalculation of saliency maps for Figure 6b-c using a different attribution method, Integrated Gradients (IG). A)** Saliency map comparison of MelanA and SOX10 models using IG. The IG method uses a different reasoning than Guided Grad-CAM to attribute salience to individual pixels, although it can be less intuitive when applied to images. Interestingly, the regions where independent IG calculations on each model converge on the same pixels (blue) qualitatively appear more coherent than those specific to either model alone (red, green). **B)** Agreement map comparing MelanA and SOX10 attributions on the same tissue region. Grayscale tissue denotes regions where the MelanA model confidence threshold is < 0.9. We only calculate attributions within the colored (high-confidence) region.

### Supplemental Tables

| WSI ID | Stain Type | Institution | # of Sections | Stain Color |
|---|---|---|---|---|
| WSI-42 | SOX10 | UCSF | 2 | Red |
| WSI-44 | SOX10 | UCSF | 2 | Red |
| WSI-47 | SOX10 | UCSF | 3 | Red |
| WSI-52 | SOX10 | Stanford | 6 | Red |
| WSI-55 | SOX10 | Stanford | 2 | Red |
| WSI-57 | SOX10 | Stanford | 1 | Brown |
| WSI-58 | SOX10 | Stanford | 1 | Brown |
| WSI-59 | SOX10 | Stanford | 4 | Brown |
| WSI-60 | SOX10 | Stanford | 2 | Red |
| WSI-61 | SOX10 | Stanford | 3 | Red |
| WSI-02 | MELA | UCSF | 4 | Red |
| WSI-03 | MELA | UCSF | 2 | Red |
| WSI-06 | MELA | UCSF | 7 | Red |
| WSI-07 | MELA | UCSF | 4 | Red |
| WSI-09 | MELA | UCSF | 1 | Red |
| WSI-12 | MELA | UCSF | 2 | Red |
| WSI-13 | MELA | UCSF | 3 | Red |
| WSI-14 | MELA | UCSF | 5 | Red |
| WSI-15 | MELA | UCSF | 1 | Red |
| WSI-16 | MELA | UCSF | 3 | Red |
| WSI-28 | MelPro | UCSF | 2 | Red |
| WSI-30 | MelPro | Stanford | 1 | Red |
| WSI-32 | MelPro | Stanford | 2 | Red |

**Supplemental Table 1.** Overview of whole slide images used to train and evaluate the models.

| WSI ID | Stain Type | Institution | Sections (HE) | Sections (IHC) | Stain Color | Preview Notes | Action | Alignment Status |
|---|---|---|---|---|---|---|---|---|
| WSI-01 | MELA | UCSF | 6 | 6 | Red | Tissue itself is strange. Previously removed | Not Used | Not Aligned |
| WSI-02 | MELA | UCSF | 8 | 8 | Red | 3rd level image damaged -- damage may impact alignment preview | Used | Aligned |
| WSI-03 | MELA | UCSF | 6 | 2 | Red | 6 slices compared to 2 rotated slices -- 3rd slice on top removed bc of bad alignment | Used | Aligned |
| WSI-04 | MELA | UCSF | 2 | 1 | Red | Good -- can't preview due to huge size disparity -- going to assume at highest res it will work out -- **Have to manually add params | Attempted | Aligned Poorly |
| WSI-05 | MELA | UCSF | 3 | 3 | Red | Iffy -- 3rd pairing is mismatch at equal sizing so removed -- other two may align, but not likely to very well | Not Used | Not Aligned |
| WSI-06 | MELA | UCSF | 8 | 8 | Red | Good | Used | Aligned |
| WSI-07 | MELA | UCSF | 5 | 5 | Red | Good -- keeping segment 1 | Used | Aligned |
| WSI-08 | MELA | UCSF | 8 | 4 | Red | IHC images on wildly larger scale than HE -- can't preview -- can't ECC align | Not Used | Not Aligned |
| WSI-09 | MELA | UCSF | 8 | 8 | Red | Good -- skip first 2 smallest segments as they are difficult to align | Used | Aligned |
| WSI-10 | MELA | UCSF | 8 | 8 | Red | Good -- remove bottom 2 segments as cannot ECC align between HE and IHC | Attempted | Aligned Poorly |
| WSI-11 | MELA | UCSF | 4 | 4 | Red | Good -- likely too big for opencv -- skip for now and use previous alignments | Attempted | Aligned Poorly |
| WSI-12 | MELA | UCSF | 4 | 2 | Red | Good | Used | Aligned |
| WSI-13 | MELA | UCSF | 12 | 12 | Red | Good -- exclude bottom 2 segments by size -- removed section 10-9 HE-IHC for bad alignment | Used | Aligned |
| WSI-14 | MELA | UCSF | 8 | 8 | Red | Good | Used | Aligned |
| WSI-15 | MELA | UCSF | 18 | 6 | Red | Good -- removed bottom slice from each trio (total of 6 slices) due to bad segmentation | Used | Aligned |
| WSI-16 | MELA | UCSF | 6 | 6 | Red | May not be able to align due to size disparity | Used | Aligned |
| WSI-17 | MELA | UCSF | 10 | 10 | Red | Good -- ignore middle 2 slices for all pairings | Attempted | Aligned Poorly |
| WSI-18 | MELA | UCSF | 6 | 2 | Red | Poor pairing -- can't preview -- likely can't align | Not Used | Not Aligned |
| WSI-19 | MELA | UCSF | 5 | 5 | Red | Good -- remove 5<->5 | Attempted | Aligned Poorly |
| WSI-20 | MELA | Stanford | 3 | 3 | Brown | Good | Not Used | Aligned |
| WSI-21 | MELA | Stanford | 3 | 3 | Brown | IHC sections are missing bottom half + angle disparity | Not Used | Not Aligned |
| WSI-22 | MELA | Stanford | 4 | 4 | Brown | IHC sections are missing bottom half + angle disparity | Not Used | Not Aligned |
| WSI-23 | MELA | Stanford | 2 | 2 | Brown | HE seg1 contains schlieren lines on right side -- Seg2 shapes different | Attempted | Aligned Poorly |
| WSI-24 | MELA | Stanford | 4 | 2 | Brown | IHC seg2 needs to be resampled | Not Used | Not Aligned |
| WSI-25 | MELA | Stanford | 1 | 1 | Brown | Bad IHC stain | Not | Not |

| | | | | | | | | |
|---|---|---|---|---|---|---|---|---|
| | | | | | | | Used | Aligned |
| WSI-26 | MELA | Stanford | 1 | 1 | Brown | Good | Attempted | Not Aligned |
| WSI-27 | MelPro | UCSF | 12 | 4 | Red | Good | Not Used | Not Aligned |
| WSI-28 | MelPro | UCSF | 18 | 6 | Red | Good -- bottom 3 IHCs are not attached to slide correctly -- applying only to top 3 | Used | Aligned |
| WSI-29 | MelPro | Stanford | 6 | 2 | Red | Good --alignments for pairs 1-3 may be difficult -- Copy in p16 | Attempted | Not Aligned |
| WSI-30 | MelPro | Stanford | 3 | 3 | Red | Good -- Slight amount of IHC bottom missing in comparison to H&E | Used | Aligned |
| WSI-31 | MelPro | Stanford | 3 | 3 | Red | Good -- IHC 2 may be treated as 2 segments instead of 1 (examine whether chroma 0. vs 1. is necessary) | Not Used | Aligned |
| WSI-32 | MelPro | Stanford | 3 | 3 | Red | Good -- NA | Used | Aligned |

**Supplemental Table 2.** MelanA and MelPro whole slide image dataset metadata and manual inspection notes.

| WSI ID | Stain Type | Institution | Sections (HE) | Sections (IHC) | Stain Color | Preview Notes | Action | Alignment Status |
|---|---|---|---|---|---|---|---|---|
| WSI-33 | SOX10 | UCSF | 3 | 2 | Black | 3 small H&E slices provided. 2 slightly larger IHC slices -- can't perform alignment preview | Not Used | Not Aligned |
| WSI-34 | SOX10 | UCSF | 2 | 1 | Black | 2 small slices compared to 1 large rotated slice -- can't perform alignment preview | Not Used | Not Aligned |
| WSI-35 | SOX10 | UCSF | 2 | 1 | Black | 2 small slices compared to 1 large rotated slice -- can't perform alignment preview | Not Used | Not Aligned |
| WSI-36 | SOX10 | UCSF | 4 | 2 | Black | 4 slices compared to 2 larger rotated slices only -- can't perform alignment preview | Not Used | Not Aligned |
| WSI-37 | SOX10 | UCSF | 2 | 2 | Black | 2 small slices compared to 1 large *broken* rotated slice -- can't perform alignment preview | Not Used | Not Aligned |
| WSI-38 | SOX10 | UCSF | 2 | 1 | Black | 2 small slices compared to 1 large *broken* slice -- can't perform alignment preview | Not Used | Not Aligned |
| WSI-39 | SOX10 | UCSF | 2 | 1 | Black | 2 small slices compared to 1 large rotated slice -- can't perform alignment preview | Not Used | Not Aligned |
| WSI-40 | SOX10 | UCSF | 3 | 2 | Black | Multiple slices compared to 2 separated portion of the same section --can't perform alignment preview | Not Used | Not Aligned |
| WSI-41 | SOX10 | UCSF | 8 | 4 | Black | Tissue is poor/choppy | Not Used | Not Aligned |
| WSI-42 | SOX10 | UCSF | 3 | 3 | Red | Good | Used | Aligned |
| WSI-43 | SOX10 | UCSF | 8 | 4 | Red | Good -- remove bottom slice tissue broken from IHC -- **WSI throws an error for align upon chroma change. Blurry IHC. | Not Used | Aligned |
| WSI-44 | SOX10 | UCSF | 6 | 2 | Red | Good -- All alignments are likely as good as they will get, but not optimal due to fundamental differences in shape between HE and IHC excisions | Used | Aligned |
| WSI-45 | SOX10 | UCSF | 3 | 3 | Red | Iffy -- most segments are too differently shaped / sized to incorporate -- May need to use original images rather than ECC | Not Used | Not Aligned |
| WSI-46 | SOX10 | UCSF | 12 | 12 | Brown | Good -- STAIN (GREY/BROWN) IS DIFFERENT THAN ORIGINAL LABEL. Negative control. | Not Used | Aligned |
| WSI-47 | SOX10 | UCSF | 6 | 6 | Red | Good -- does not perform well on ECC alignment | Used | Aligned |
| WSI-48 | SOX10 | UCSF | 12 | 4 | Red | Poor pairing and can't tell if slice is worthwhile | Not Used | Not Aligned |
| WSI-49 | SOX10 | UCSF | 8 | 8 | Red | Good | Attempted | Aligned Poorly |
| WSI-50 | SOX10 | UCSF | 6 | 6 | Red | Good -- stain somewhat light | Attempted | Aligned Poorly |
| WSI-51 | SOX10 | Stanford | 3 | 3 | Brown | Spurious -- Slices are huge and IHC stain is hard to separate from BG -- make sure masks are correctly matched | Attempted | Aligned Poorly |
| WSI-52 | SOX10 | Stanford | 3 | 3 | Red | COPY of slices in MELA -- HE images seem like bigger area | Used | Aligned |
| WSI-53 | SOX10 | Stanford | 4 | 3 | Red | COPY of slices in MELA -- Could be hard to align if we don't fix angle disparity | Attempted | Aligned Poorly |
| WSI-54 | SOX10 | Stanford | 4 | 4 | Brown | Good | Attempte | Aligned |

| | | | | | | | | |
|---|---|---|---|---|---|---|---|---|
| | | | | | | | d | Poorly |
| WSI-55 | SOX10 | Stanford | 4 | 4 | Red | Good | Used | Aligned |
| WSI-56 | SOX10 | Stanford | 9 | 3 | Red | Good | Attempted | Aligned Poorly |
| WSI-57 | SOX10 | Stanford | 1 | 1 | Brown | Good -- See if difference in angle causes alignment to fail | Used | Aligned |
| WSI-58 | SOX10 | Stanford | 1 | 1 | Brown | Good | Used | Aligned |
| WSI-59 | SOX10 | Stanford | 4 | 4 | Brown | Good | Used | Aligned |
| WSI-60 | SOX10 | Stanford | 3 | 3 | Red | Good | Used | Aligned |
| WSI-61 | SOX10 | Stanford | 3 | 3 | Red | Good - Yellow smudge may cause problem with matches… make sure they are correct | Used | Aligned |

**Supplemental Table 3.** SOX10 whole slide image dataset metadata and manual inspection notes.

| TCGA WSI SVS Filename | Tissue Source |
|---|---|
| TCGA-22-4594-01Z-00-DX1.3FCEBC89-8473-4841-87A2-F84AF58A7793.svs | Lung Squamous Cell Carcinoma |
| TCGA-2J-AABA-01Z-00-DX1.93B2B4EF-C302-4D00-ABE3-4862ACC81659.svs | Pancreatic Adenocarcinoma |
| TCGA-2J-AABK-01Z-00-DX1.AF5DE1FD-40EE-4149-8918-B53EC2DF727E.svs | Pancreatic Adenocarcinoma |
| TCGA-3A-A9I7-01Z-00-DX1.23EE4A93-A298-4522-837E-3EE10172D66C.svs | Pancreatic Adenocarcinoma |
| TCGA-3A-A9IB-01Z-00-DX1.77855A18-9E12-4F6A-8FBB-B5057656C493.svs | Pancreatic Adenocarcinoma |
| TCGA-3A-A9IH-01Z-00-DX1.578316D1-186E-4AE4-BD6A-DA426DE87829.svs | Pancreatic Adenocarcinoma |
| TCGA-3A-A9IL-01Z-00-DX1.BEB57CA5-223D-4330-BFFF-8202DCC857F3.svs | Pancreatic Adenocarcinoma |
| TCGA-3A-A9IN-01Z-00-DX1.A4FED037-D993-4F71-B422-14FC4E468B4C.svs | Pancreatic Adenocarcinoma |
| TCGA-3A-A9J0-01Z-00-DX1.322C8475-A1E3-4877-B3B5-921FDDB9698F.svs | Pancreatic Adenocarcinoma |
| TCGA-43-A56V-01Z-00-DX1.AA93FE03-FA7D-42C4-A118-B98C2400D9DA.svs | Lung Squamous Cell Carcinoma |
| TCGA-60-2722-01Z-00-DX1.f3781266-e8dc-4386-9702-5b29e6f2cfa3.svs | Lung Squamous Cell Carcinoma |
| TCGA-66-2742-01Z-00-DX1.8fdd6990-a08c-457b-80e4-586c619a784e.svs | Lung Squamous Cell Carcinoma |
| TCGA-A1-A0SE-01Z-00-DX1.04B09232-C6C4-46EF-AA2C-41D078D0A80A.svs | Breast Invasive Carcinoma |
| TCGA-A2-A04U-01Z-00-DX1.06D17357-46A8-4DC3-A22B-2F4EB6EE3F79.svs | Breast Invasive Carcinoma |
| TCGA-A2-A0CZ-01Z-00-DX1.A433A414-4F1B-4F99-8FD9-E64803F5E042.svs | Breast Invasive Carcinoma |
| TCGA-A8-A09R-01Z-00-DX1.392580F3-0CE5-4EDB-91CF-814AAD0DB649.svs | Breast Invasive Carcinoma |
| TCGA-AO-A0JC-01Z-00-DX1.C8DD421B-9799-4FE7-9224-5EAC6ED1028E.svs | Breast Invasive Carcinoma |
| TCGA-AO-A1KQ-01Z-00-DX1.CAB7D9A5-7030-4A33-BE51-9B04D67A7676.svs | Breast Invasive Carcinoma |
| TCGA-AR-A1AX-01Z-00-DX1.2389D54F-545E-499E-B392-DD731834460A.svs | Breast Invasive Carcinoma |
| TCGA-BH-A0DI-01Z-00-DX1.6A42D535-8842-4C36-8299-A40E9E56759D.svs | Breast Invasive Carcinoma |
| TCGA-D8-A1XS-01Z-00-DX2.ED8BBDB4-CEA6-4E47-8214-4666F3CC6E44.svs | Breast Invasive Carcinoma |
| TCGA-E9-A22D-01Z-00-DX1.b2867437-0add-4b7d-8002-fb09ed961942.svs | Breast Invasive Carcinoma |
| TCGA-F2-A44H-01Z-00-DX1.98C75E19-10DE-434A-AF1B-CDD182F6EDD5.svs | Pancreatic Adenocarcinoma |
| TCGA-F2-A7TX-01Z-00-DX1.2FB4B966-3F76-4BB7-B1E8-D6F651665479.svs | Pancreatic Adenocarcinoma |
| TCGA-FB-A4P5-01Z-00-DX1.D5440110-D217-4B4C-A8D2-7261B430F440.svs | Pancreatic Adenocarcinoma |
| TCGA-FB-A78T-01Z-00-DX1.1DC04A89-2428-489B-A70E-0D9C6D2A5E61.svs | Pancreatic Adenocarcinoma |
| TCGA-H6-A45N-01Z-00-DX1.80D3E1A9-02EB-4897-9632-F6FC00B3FA0F.svs | Pancreatic Adenocarcinoma |
| TCGA-HN-A2OB-01Z-00-DX1.14F1FBFB-4540-43CE-9D79-5BC628640424.svs | Breast Invasive Carcinoma |
| TCGA-HV-A5A4-01Z-00-DX1.00C72860-A4C4-41FB-87BA-7C4381FAF2BD.svs | Pancreatic Adenocarcinoma |
| TCGA-HZ-8005-01Z-00-DX1.e49bbccf-eab2-4f2f-b882-406b90fb2020.svs | Pancreatic Adenocarcinoma |
| TCGA-HZ-8315-01Z-00-DX1.F6B3F80E-3630-426E-AB2C-7F2EC5B63BFC.svs | Pancreatic Adenocarcinoma |
| TCGA-HZ-8317-01Z-00-DX1.BD28612C-D35D-4664-8B88-A85EF99013AB.svs | Pancreatic Adenocarcinoma |
| TCGA-HZ-8317-01Z-00-DX2.FDB366FF-AAA9-4FDC-A1F4-BA021904ED94.svs | Pancreatic Adenocarcinoma |
| TCGA-HZ-8637-01Z-00-DX1.5943021F-C94B-4CED-B45F-7A288F7188E0.svs | Pancreatic Adenocarcinoma |

| | |
|---|---|
| TCGA-HZ-8638-01Z-00-DX1.AD9F30CA-8943-493E-8603-7D1CF41056E6.svs | Pancreatic Adenocarcinoma |
| TCGA-HZ-A77O-01Z-00-DX1.C0F88C8F-C68C-457B-A0FF-3B483FCE7385.svs | Pancreatic Adenocarcinoma |
| TCGA-IB-7644-01Z-00-DX1.A2E77093-90D2-4ED3-90EB-F14A03C3DA57.svs | Pancreatic Adenocarcinoma |
| TCGA-IB-8127-01Z-00-DX1.C7035E56-9D24-4EEA-A09E-8276382193CC.svs | Pancreatic Adenocarcinoma |
| TCGA-IB-A5SS-01Z-00-DX1.899575C7-D239-4A04-8827-044F0D8868C8.svs | Pancreatic Adenocarcinoma |
| TCGA-IB-AAUV-01Z-00-DX1.045691CD-E0F8-4992-BF49-43AF7F83C97A.svs | Pancreatic Adenocarcinoma |
| TCGA-NC-A5HT-01Z-00-DX1.9295B0E3-37FE-4914-AFB3-78B56C893B6D.svs | Lung Squamous Cell Carcinoma |
| TCGA-OL-A97C-01Z-00-DX1.BDEEDEE2-6D07-4046-A8A9-D6FF8F337393.svs | Breast Invasive Carcinoma |
| TCGA-US-A774-01Z-00-DX1.522FF138-153F-488A-BBBC-5EA68EFD80C7.svs | Pancreatic Adenocarcinoma |
| TCGA-UU-A93S-01Z-00-DX1.C4809779-DF5F-4F5D-A78C-B7F95F2D050F.svs | Breast Invasive Carcinoma |
| TCGA-Z5-AAPL-01Z-00-DX1.30371C08-9075-44A9-8ED7-560256D65A7C.svs | Pancreatic Adenocarcinoma |

**Supplemental Table 4.** TCGA identifiers for SVS whole slide images and tissue sources for negative-control TCGA non-skin H&E tiles used in model training.

| Tissue Source | WSI Count |
|---|---|
| Pancreatic Adenocarcinoma | 27 |
| Breast Invasive Carcinoma | 13 |
| Lung Squamous Cell Carcinoma | 5 |

**Supplemental Table 5.** Summary of tissue sources for negative-control TCGA non-skin H&E tiles used in model training.

| Fold | Train and Validation | | Test | |
|---|---|---|---|---|
| | Positive | Negative | Positive | Negative |
| MelanA 1 | 16,214 | 355,753 | 1,025 | 20,924 |
| MelanA 2 | 15,034 | 354,610 | 800 | 9,422 |
| MelanA 3 | 13,102 | 317,536 | 1,225 | 20,614 |
| MelanA 4 | 15,281 | 383,511 | 855 | 12,648 |
| MelanA 5 | 12,712 | 329,430 | 1,505 | 16,186 |
| Sox10 1 | 8,788 | 282,041 | 1,099 | 20,507 |
| Sox10 2 | 8,758 | 333,967 | 958 | 8,926 |
| Sox10 3 | 8,141 | 283,539 | 489 | 14,778 |
| Sox10 4 | 10,278 | 283,860 | 407 | 5,454 |
| Sox10 5 | 8,432 | 269,898 | 389 | 8,517 |

**Supplemental Table 6.** Breakdown of positive and negative tile counts for each fold.
